# Proteomic and lipidomic profiling of immune cell-derived subpopulations of extracellular vesicles

**DOI:** 10.1101/2025.05.08.652936

**Authors:** Anna Lischnig, Nasibeh Karimi, Per Larsson, Karin Ekström, Rossella Crescitelli, Anna-Carin Olin, Cecilia Lässer

**Affiliations:** Krefting Research Centre, Department of Internal Medicine and Clinical Nutrition, Institute of Medicine, Sahlgrenska Academy, University of Gothenburg, Gothenburg, Sweden; Chalmers Mass Spectrometry Infrastructure, Department of Life Sciences, Chalmers University of Technology, Gothenburg, Sweden; Occupational and Environmental Medicine, Department of Public Health and Community Medicine, Institute of Medicine, Sahlgrenska Academy, University of Gothenburg, Gothenburg, Sweden; Sahlgrenska Center for Cancer Research, Department of Surgery, Institute of Clinical Sciences, Sahlgrenska Academy, University of Gothenburg, Gothenburg, Sweden; Department of Surgery, Sahlgrenska University Hospital, Gothenburg, Sweden

**Keywords:** Exosomes, Microvesicles, Extracellular Vesicles, Protein cargo, Lipid composition

## Abstract

**Introduction:** Extracellular vesicles (EVs) are a heterogeneous group of membrane-enclosed vesicles released by cells. They play important roles in intercellular communication and contribute to several physiological and pathological processes. Cells release subpopulations of EVs with distinct biogenesis and functions, however, we currently have few markers to differentiate them. This study, therefore, aimed to determine proteomic and lipidomic differences among four EV subpopulations of varying sizes and densities.

**Methods:** Large and small EVs (L-EVs and S-EVs) were isolated from two immune cell lines by differential ultracentrifugation at 16,500 × g and 118,000 × g, respectively. The crude EVs were then further separated by density cushion centrifugation. EVs were isolated from the interphase between 1.079-1.146 and 1.146-1.185 g/ml, hereafter referred to as low density (LD) and high density (HD), respectively. This resulted in four subpopulations of EVs: L-EVs LD, L-EVs HD, S-EVs LD, and S-EVs HD. The morphology, size, and yield of EVs were determined by nanoparticle tracking analysis, electron microscopy, and western blot. The proteome and lipidome of the four subpopulations of EVs were determined with mass spectrometry.

**Results:** A total of 5364 proteins were quantified in the dataset. L-EV and S-EVs as well as LD and HD were well separated. Briefly, L-EVs LD were enriched in mitochondrial proteins such as the TIMM/TOMM complex, MICOS, and ATP5 proteins. In contrast, L-EVs HD were enriched in proteins associated with the cytoskeleton, such as KIF proteins. Furthermore, S-EVs LD were enriched in tetraspanins and ESCRT machinery proteins, while S-EVs HD were enriched in histones, CCT proteins, and proteins from the complement pathway. Proteins such as flotillins, RABs, annexins, and integrins were enriched in two or several subpopulations. Furthermore, 107 lipids were quantified, and the most abundant lipids in EVs were phosphatidylcholine (PC), sphingomyelin, and phosphatidylethanolamine (PE). The most profound difference was that PE was less abundant in L-EVs LD as compared to the other EV subtypes, and ceramides were enriched in L-EVs as compared to S-EVs.

**Conclusion:** This study demonstrates that the proteome and lipidome differ in EV subpopulations separated based on size and density. Furthermore, it validates several protein groups that have previously been suggested to be enriched in either S-EVs or L-EVs.

## 1. Introduction

Extracellular vesicles (EVs) are small lipid membrane-bound particles released by cells that play critical roles in intercellular communication by transporting proteins, lipids, and nucleic acids between cells (1). EVs have been shown to play a role in a variety of biological processes including homeostasis, inflammation, neurodegenerative diseases, and cancer (2, 3). In addition, EVs are studied for their clinical use as biomarkers and therapeutic vehicles for several diseases (4, 5). Large-scale studies of EV composition and cargo are therefore pivotal for understanding the complex biological functions and mechanisms of EVs.

EVs is an umbrella term for several different subpopulations of vesicles, including, but not limited to exosomes, microvesicles, and apoptotic bodies, which differ in size, biogenesis, and function (6). However, their nomenclature remains inconsistent, and most isolation protocols cannot distinguish EVs based on biogenesis, yet those terms are still applied. Therefore, we used the terms large EVs (L-EVs) for EVs isolated at 10,000-20,000 × g and the term small EVs (S-EVs) for vesicles isolated at centrifugation forces above 100,000 × g to describe the EV subpopulations of this study and the studies we reference. While several proteomics studies have examined the protein content of S-EVs, less is known about L-EVs. Comparisons of multiple EV subpopulations within the same study are even rarer. This has led to a knowledge gap regarding subpopulation-specific proteins. The characterization of the content and cargo of EV subpopulations is crucial for advancing EV research across a variety of applications. These markers can enhance our understanding of EV biology, aid in the discovery of biomarkers, and improve the characterization of EV isolation and purification processes.

Lipids not only serve as structural components of cell and vesicle membranes and as energy stores but also function as signalling molecules (7). Although the lipid content of EVs is as important as their protein content for understanding biogenesis and function, few comprehensive lipidomic EV subpopulation studies have been conducted. The aim of this study is therefore to determine and compare the proteome and lipid composition of four subpopulations of EVs. Undertaking this will allow us to determine which proteins and lipids are uniquely expressed in certain subpopulations and which are common among EV subtypes.

Here we present an in-depth analysis of the proteomes and lipidomes of four EV subpopulations with different sedimentation properties. Four different sample types, namely L-EVs low density (LD), L-EVs high density (HD), S-EVs LD, and S-EVs HD, were isolated using a combination of differential ultracentrifugation and subsequent density cushion centrifugation. Then we characterised the EV subtypes regarding their purity, morphology, yield, and cargo. The analysis of the proteome by TMT-LC-MS/MS revealed the differential expression of several proteins and protein groups in the different EV subtypes. The analysis of the lipidomes showed differences in the relative abundance of lipid classes and single lipids.

## 2. Material and methods

### 2.1 Cell cultures

The mast cell line HMC-1 (8, 9) was cultured in Iscove’s Modified Dulbecco’s Medium (IMDM; Cytiva, HyClone laboratories, Inc., and Gibco, Thermo Fisher Scientific Inc.) supplemented with 1.2 mM 1-Thioglycerol (Sigma-Aldrich). The monocyte cell line THP-1 (10) was cultured in RPMI 1640 Medium (Cytiva) supplemented with 0.05 mM 2-Mercaptoethanol (Gibco). Additionally, all media were supplemented with 10% EV-depleted foetal bovine serum (FBS; Sigma-Aldrich), 100 units/mL Penicillin (Cytiva), 100 μg/mL Streptomycin (Cytiva) and 2 mM L-Glutamine (Cytiva). The FBS was depleted of EVs by ultracentrifugation at 118,000 × g_avg_ with a Type 45 Ti fixed-angle rotor (k-factor 178; 38 800 rpm, Beckman Coulter) for 18 h at 4°C. The EV-depleted FBS was sterile filtered through a 0.22 μm filter (Sarstedt AG & Co.) before being added to the media. The incubator was humidified and set to 37°C with 5% CO_2_. Cells were seeded at 5 × 10^5^ cells/ml, and the cells were cultured for 3 - 4 days before EVs were isolated. For each isolation, approximately 600 ml of conditioned cell culture medium was harvested.

### 2.2. Enrichment of large and small EVs by differential ultracentrifugation

EVs were isolated from HMC-1 and THP-1 conditioned cell culture medium by differential ultracentrifugation (**Figure 1A**). All centrifugation steps were performed at 4°C. First, the medium was sequentially centrifuged for 20 minutes at 300 × g and 20 minutes at 2,000 × g to pellet the cells, cell debris and apoptotic bodies. Next, the supernatant was centrifuged for 20 minutes at 16,500 × g_avg_ with a Type 45 Ti fixed angle rotor (k-factor 1277; 14,500 rpm), hereafter, this pellet is referred to as crude L-EVs. The supernatant was then used for the final ultracentrifugation at 118,000 × g_avg_ for 2.5 h with a Type 45 Ti fixed angle rotor (k-factor 178; 38,800 rpm); hereafter, this pellet is referred to as crude S-EVs. All pellets were resuspended with phosphate-buffered saline (PBS) and stored at −80°C.

**Figure 1.**
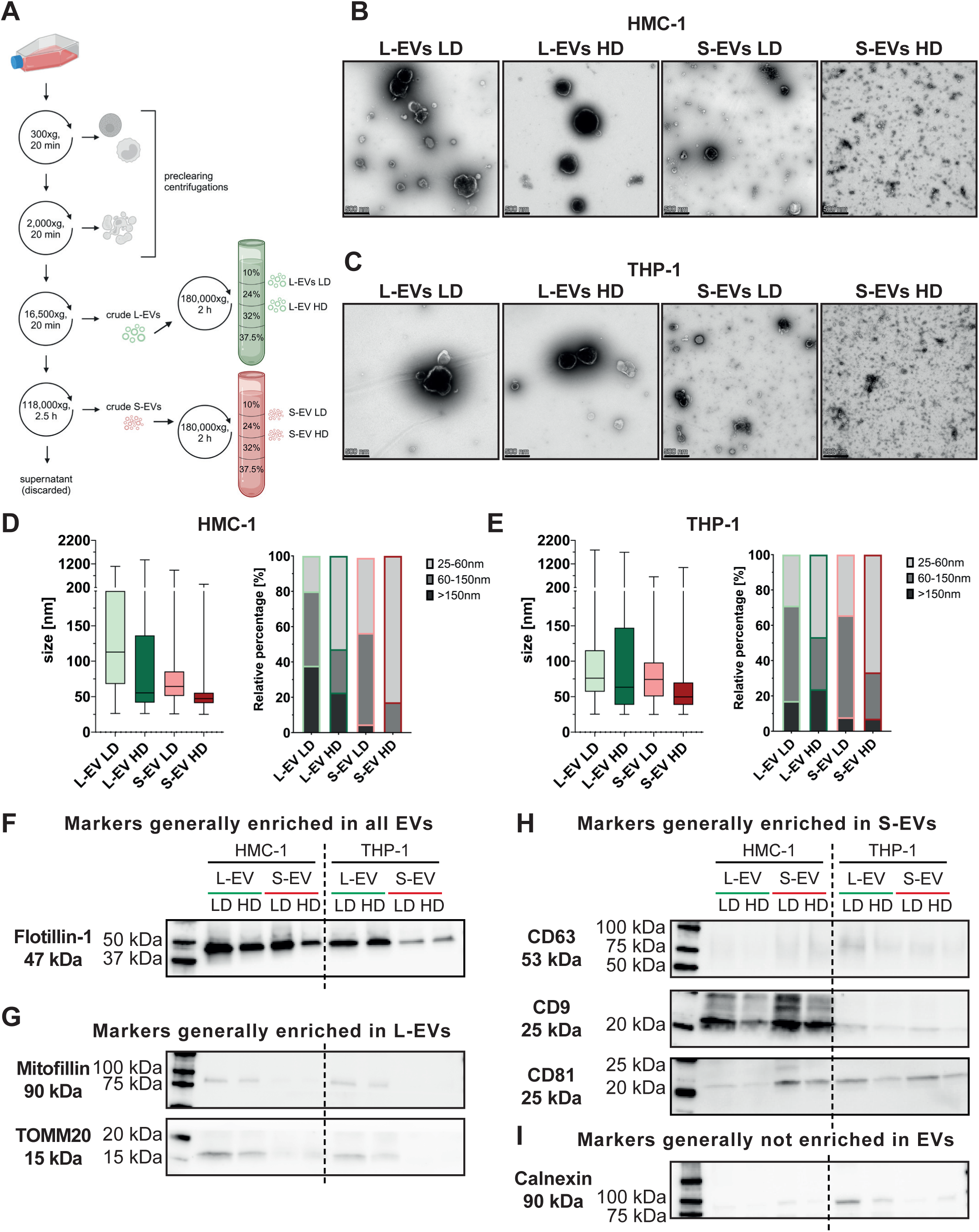
Characterization of four subpopulations of EVs isolated from immune cells. **(A)** Schematic overview of the isolation protocol including both differential ultracentrifugation and a density cushion. **(B-C)** Five micrograms of vesicles were loaded onto grids, negative stained, and evaluated by transmission electron microscopy. Representative micrographs are shown from HMC-1 (B) and THP-1 (C). The scale bars represent 500 nm. **(D-E)** For each vesicle subtype, 41-45 micrographs were acquired. All structures in the micrographs were manually measured (**Table 3**). In total, the diameters of over 8,500 and 5,000 structures were measured in HMC-1 (D) and THP-1 (E), respectively. Two biological replicates were analysed per EV type. **(F-I)** Western blot was used to investigate the presence of markers generally enriched in all EVs (F), markers generally enriched in L-EVs (G), markers generally enriched in S-EVs (H), and markers generally not enriched in EV (I). A total of 6µg was loaded per sample. HD, high density; LD, low density; L-EVs, large extracellular vesicles; S-EVs small extracellular vesicles

### 2.3. Enrichment of low-density and high-density EVs by density cushion centrifugation

The crude L-EVs and S-EVs were purified on bottom-loaded density cushions and EVs at two different densities were collected (**Figure 1A** and **Supplementary** Figure 1). The densities at which the EV subtypes were isolated were chosen based on previous experiences (4, 11, 12). Briefly, the crude EV samples of two to three crude EV isolations were pooled, and PBS was added to reach a total volume of 1.5 ml. This was then mixed with 2.5 ml 60% iodixanol (OptiPrep, Merck). The 4 ml sample/iodixanol mix with an iodixanol concentration of 37.5% was loaded at the bottom of the tube (Open-Top Thinwall Ultra-Clear Tube, 13.2 ml, 14 x 89 mm, Beckman Coulter). Then 2.65 ml of 32%, 2.65 ml of 24%, followed by 2.65 ml of 10% iodixanol solution, was layered on top. The cushion was centrifuged at 180,000 × g_avg_ (SW 41 Ti rotor, k-factor 145, 38 000 rpm) for 2 h at 4°C. After the centrifugation, the interphases between the 10% and 24% iodixanol layer (1.079 −1.146 g/ml) and the interphase between the 24% and 32% iodixanol layer (1.146 - 1.185 g/ml) were collected, these fractions are referred to as LD and HD fractions hereafter, respectively.

### 2.4. Characterization of EVs

#### 2.4.1. Particle and protein measurements

Particle concentrations were determined on a ZetaView PMX instrument (Particle Metrix, Germany). Auto alignment and daily performance routines were performed according to the manufacturer’s instructions. Briefly, the ZetaView Particle Size Standard Polystyrene 100 nm (Applied Microspheres, Leusden, The Netherlands) was diluted to 1:250,000 immediately before use and calibration was performed. The EV samples were diluted 1000-50,000-fold in PBS immediately prior to the acquisition. Acquisition settings, with a sensitivity of 80 and a shutter speed of 100, were maintained for all samples. Eleven positions with two cycles were recorded for each sample. Analysis parameters were consistent across all samples, with a minimum size of 5, a maximum size of 1000, and a minimum brightness of 20. The software ZetaView was used for the analysis (Version 8.05.12 SP1).

The Pierce™ BCA Protein Assay Kit (Thermo Scientific) was used according to the manufacturer’s instructions. Samples were measured in Clear Flat-Bottom Immuno Nonsterile 96-Well Plates (Thermo Fisher Scientific) using a Varioskan LUX microplate reader (SkanIt Software 4.1 for Microplate Readers RE, ver. 4.1.0.43).

#### 2.4.2. Western blot

The EV samples were thawed and diluted, with 6 µg of protein loaded for all samples. Subsequently, 4 × Laemmli buffer (Bio-Rad Laboratories) was added to the samples. For samples requiring reducing conditions, 100 µL of β-mercaptoethanol was added to 900 µL of 4 × Laemmli buffer. The samples were denatured at 95°C for 5 minutes before loading onto Mini-Protean TGX precast 4 to 20% gels (Bio-Rad Laboratories). As molecular weight marker, 4 µL Precision Plus Protein™ WesternC™ Blotting Standard (Bio-Rad Laboratories) was used. The proteins were separated at 180 V for about 45 minutes. Blotting onto PVDF membranes was performed on the Trans-Blot Turbo Transfer system (Bio-Rad Laboratories) set to 1.3 A and 25 V for 7 minutes using transfer buffer (100 mL Trans-Blot Turbo stock solution (Bio-Rad Laboratories), 100 mL Ethanol, and 300 mL distilled water). The membranes were blocked with EveryBlot Blocking Buffer (Bio-Rad Laboratories) for 5 minutes and then incubated with the primary antibodies (**Table 1**) diluted in EveryBlot Blocking Buffer overnight at 4°C.

**Table 1.** Primary antibodies used for western blot.

| **Protein** | **Clone** | **Number** | **Supplier** | **Host** | **Condition** | **Dilution** |
| --- | --- | --- | --- | --- | --- | --- |
| **CD81** | M38 | ab79559 | Abcam | mouse | not reducing | 1:1000 |
| **CD63** | H5C6 | 556019 | BD Pharmingen | mouse | not reducing | 1:1000 |
| **CD9** | MM2/57 | CBL162 | EMD Millipore | mouse | not reducing | 1:1000 |
| **Calnexin** | C5C9 | 2679 | Cell Signalling Technology | rabbit | reducing | 1:1000 |
| **Flotillin-1** | EPR6041 | ab133497 | Abcam | rabbit | reducing | 1:1000 |
| **Mitofilin** | EPR8749 | ab137057 | Abcam | rabbit | reducing | 1:1000 |
| **TOMM20** | EPR15581-54 | ab186735 | Abcam | rabbit | reducing | 1:2000 |

After primary antibody incubation, the membranes were washed three times with TBS (Bio-Rad Laboratories) supplemented with 0.05% Tween 20 (TBS/Tween 20). Membranes were incubated with the secondary antibodies (**Table 2**) and Precision Protein StrepTactin-HRP Conjugate (1:10 000; Bio-Rad Laboratories) diluted in EveryBlot Blocking Buffer at RT for one hour. After incubation, the membranes were washed four times in TBS/Tween 20. The blots were imaged with SuperSignal West Femto Maximum Sensitivity Substrate (Thermo Fisher Scientific) and the ChemiDoc Imaging System (Bio-Rad Laboratories). The secondary antibodies and StrepTactin-HRP were evaluated for unspecific binding to the samples in reducing and non-reducing conditions.

**Table 2.** Secondary antibodies used for western blot.

| **Antibody** | **Clonality** | **Number** | **Supplier** | **Host** | **Dilution** |
| --- | --- | --- | --- | --- | --- |
| anti-rabbit<br>IgG | polyclonal | NA9340V | GE Healthcare UK<br>Limited | donkey | 1:5000 |
| anti-mouse<br>IgG | polyclonal | NA9310V | GE Healthcare UK<br>Limited | sheep | 1:5000 |

**Table 3.** Number of vesicles measured with TEM.

|  | **HMC-1** |  |  |  | **THP-1** |  |  |  |
| --- | --- | --- | --- | --- | --- | --- | --- | --- |
| **EV type** | **L-EVs**<br>**LD** | **L-EVs**<br>**HD** | **S-EVs**<br>**LD** | **S-EVs**<br>**HD** | **L-EVs**<br>**LD** | **L-EVs**<br>**HD** | **S-EVs**<br>**LD** | **S-EVs**<br>**HD** |
| **Number**<br>**of**<br>**images** | 44 | 44 | 45 | 44 | 43 | 42 | 45 | 41 |
| **Number**<br>**of EVs** | 1263 | 852 | 4044 | 2358 | 1516 | 664 | 1615 | 1279 |

#### 2.4.3. Transmission electron microscopy

For negative staining, formvar/carbon-coated nickel grids (Ted Pella, Inc.) were glow discharged prior to incubation with 5 µg of EVs for 5 minutes. Grids were washed two times with H_2_O. Samples were then fixed with 2% glutaraldehyde for 10 minutes before being negative stained with 2% uranyl acetate for 1.5 minutes. The grids were then dried well before being acquired using a Talos L120C transmission electron microscope (Thermo Fisher Scientific) at 120 kV with a CCD camera.

For each of the sample types, two biological replicates were prepared on grids for imaging. Approximately a total of 22 images were acquired from three different positions on each grid. EVs were counted, and the longest diameter of each EV was measured using Fiji ImageJ 2.14.0/1.54f (37). Sample preparation, image acquisition, and measuring of EVs were conducted in a blinded manner. That is the operator who acquired the images and measured the EVs was blinded to which sample was which. This was to avoid bias towards capturing pictures or measuring vesicles of a certain size, depending on knowing which sample was analysed. The total amount of EVs measured for each EV subtype is listed in **Table 3**.

### 2.5. Proteomics

#### 2.5.1. Sample generation

For tandem mass tag (TMT)–liquid chromatography (LC)–tandem mass spectrometry (MS/MS), three independent biological replicates were used for each of the four sample types resulting in twelve samples per cell line and 24 samples in total (n=3 for each vesicle type and cell line). The TMT method used allowed the comparison of up to sixteen samples in one set. Therefore, the twelve samples for each cell line were run on two different sets. Proteomic analysis was performed at the Proteomics Core Facility at Sahlgrenska Academy, University of Gothenburg, Sweden.

#### 2.5.2. Protein digestion and labelling

Aliquots containing 40 μg of each sample (except 35 µg protein for HMC-1 L-EVs HD-2), and two reference pools made up of aliquots from each sample, were digested with trypsin using the filter-aided sample preparation (FASP) method (38). Briefly, the samples were reduced with 100 mM dithiothreitol at 56°C for 30 minutes. Samples were diluted to 1:3 with 8 M urea in 50 mM triethylammonium bicarbonate (TEAB), transferred onto Microcon-30kDa Centrifugal Filter Units (Merck), and washed four times by adding 200 µL of 8 M urea in 50 mM TEAB, two times with digestion buffer (0.5% sodium deoxycholate (SDC), 50 mM TEAB) and subsequent centrifugation at 10 000 x g. Free cysteine residues were modified using 10 mM methyl methanethiosulfonate (MMTS) solution in digestion buffer for 20 minutes at RT and the filters were then repeatedly washed with 100 µL of digestion buffer. Pierce trypsin protease (MS Grade, Thermo Fisher Scientific) in digestion buffer was added at a ratio of 1:100 relative to total protein mass and the samples were incubated at 37°C overnight. Next morning, another portion of trypsin (1:100) was added and the mixture was incubated at 37°C for 4 h. The peptides were collected by centrifugation and labelled using TMTpro 16-plex isobaric mass tagging reagents (Thermo Scientific) according to the manufacturer’s instructions. Samples from each cell line were labelled in separate sets and combined into two sets together with one reference sample. Acetonitrile was evaporated using vacuum centrifugation and SDC was removed by acidification with 10% TFA with subsequent centrifugation. The pooled samples were purified using High Protein and Peptide Recovery Detergent Removal Spin Column (Thermo Fisher Scientific) according to the manufacturer instructions, followed by Pierce Peptide Desalting Spin Columns (Thermo Scientific), following the manufacturer’s instructions.

The combined labelled sample was fractionated into 34 primary fractions by basic reversed-phase chromatography (bRP-LC) using a Dionex Ultimate 3000 UPLC system (Thermo Fischer Scientific). Peptide separations were performed on a reversed-phase XBridge BEH C18 column (3.5 μm, 3.0×150 mm, Waters Corporation) using a gradient from 8% to 40% solvent B over 48 minutes, 40% to 50% over 10 minutes followed by an increase to 100% B over 5 minutes, at flow of 400 µL/minute. Solvent A was 10 mM ammonium formate buffer at pH 10.00 and solvent B was 90% acetonitrile, 10% 10 mM ammonium formate at pH 10.00. The primary fractions were concatenated into final 17 fractions (1+18, 2+19, … 17+34), evaporated, and reconstituted in 15 µL of 3% acetonitrile, 0.2% trifluoroacetic acid.

#### 2.5.3. LC-MS/MS analysis

Each fraction was analysed on Orbitrap Fusion Tribrid mass spectrometer interfaced with Easy-nLC 1200 nanoflow liquid chromatography system (both - Thermo Fisher Scientific). Peptides were trapped on the Acclaim Pepmap 100 C18 trap column (75 μm X 2 cm, particle size 5 μm, Thermo Fischer Scientific) and separated on the in-house packed C18 analytical column (75 μm X 37 cm, particle size 3 μm) using the gradient from 5% to 12% B over 5 minutes, 12% to 35% B over 70 minutes followed by an increase to 100% B for 5 minutes, and 100% B for 10 minutes at a flow of 300 nL/minute. Solvent A was 0.2% formic acid and solvent B was 80% acetonitrile, 0.2% formic acid. Precursor ion mass spectra were recorded at 120 000 resolution, the most intense precursor ions were selected (‘top speed’ setting with a duty cycle of 3 s), fragmented using CID at collision energy setting of 30, spectra and the MS/MS spectra were recorded in ion trap with the maximum injection time of 50 ms and the isolation window of 0.7 Da. Charge states 2 to 7 were selected for fragmentation, dynamic exclusion was set to 45 s with 10 ppm tolerance. MS3 spectra for reporter ion quantitation were recorded at 50,000 resolutions with HCD fragmentation at a collision energy of 55 using the synchronous precursor selection of the 10 most abundant MS/MS fragments, with the maximum injection time of 120 ms.

#### 2.5.4. Database search and quantification

Identification and relative quantification were performed of the combined injections using Proteome Discoverer version 2.4 (Thermo Fisher Scientific). The database search was performed using the Mascot search engine v. 2.5.1 (Matrix Science, London, UK) against the Swiss-Prot human database. Trypsin was used as a cleavage rule with no missed cleavages allowed; methylthiolation on cysteine residues, TMTpro at peptide N-termini, and on lysine side chains were set as static modifications and oxidation on methionine was set as a dynamic modification. Precursor mass tolerance was set at 5 ppm and fragment ion tolerance at 0.6 Da. A percolator was used for the peptide-spectrum match (PSM) validation with a strict false discovery rate (FDR) threshold of 1%. Quantification was performed in Proteome Discoverer 2.4. The TMT reporter ions were identified with 3 mmu mass tolerance in the MS3 HCD spectra and the TMT reporter S/N values for each sample were normalized within Proteome Discoverer 2.4 on the total peptide amount. Only the quantitative results for the unique peptide sequences with the minimum SPS match % of 50 and the average S/N above 10 were taken into account for the protein quantification.

#### 2.5.5. Statistics and bioinformatics

For the analysis of the quantified proteins, the significance was calculated by paired Student’s t-test on logged values. Subsequently, GraphPad Prism 10 was used to create the volcano plots. Qlucore Omics Explorer (Qlucore Omics Explorer Version 3.9) was used for principal component analysis. After identifying the uniquely enriched proteins in the different EV subtypes the Database for Annotation, Visualization and Integrated Discovery (DAVID; https://david.ncifcrf.gov/home.jsp [accessed: 22–08–2024]) was used to determine the most enriched cellular compartments associated with these proteins.

The proteins from the five MISEV categories were created based on Table 3 in the MISEV2023 guidelines (13) and analysed and visualized with Qlucore Omics Explorer (Qlucore Omics Explorer Version 3.9) and multigroup (ANOVA) comparisons displayed as hierarchical clustering heat maps. Category 1 contained 64 proteins in our data set and the p-value was set to 0.01 for both cell lines. This generated q-values of 0.018771 and 0.01367 for HMC-1 and THP-1, respectively. The heatmaps then contained 32 and 40 proteins in HMC-1 and THP-1, respectively. Category 2 contained 44 proteins in our data set and the p-value was set to 0.01 for both cell lines. This generated q-values of 0.01843 and 0.01023 for HMC-1 and THP-1, respectively. The heatmaps then contained 22 and 36 proteins in HMC-1 and THP-1, respectively. Category 3 contained 38 proteins in our data set and the p-value was set to 0.01 for both cell lines. This generated q-values of 0.017891 and 0.014379 for HMC-1 and THP-1, respectively. The heatmaps then contained 14 and 21 proteins in HMC-1 and THP-1, respectively. Category 4 contained 19 proteins in our data set and the p-value was set to 0.01 for both cell lines. This generated q-values of 0.000503 and 0,0060883 for HMC-1 and THP-1, respectively. The heatmaps then contained 13 and 19 proteins in HMC-1 and THP-1, respectively. Category 5 contained 30 proteins in our data set and the p-value was set to 0.01 for both cell lines. This generated q-values of 0,0095448 and 0,0084668 for HMC-1 and THP-1, respectively. The heatmaps then contained 15 and 22 proteins in HMC-1 and THP-1, respectively.

The list of common EV proteins where created based on Top100 proteins in VesiclePedia (accessed: 27-08-2024), Top100 proteins in EVpedia (accessed 27-08-2024), subpopulation markers suggested by Lim *et al* in a recent paper (14), and marker for L-EVs from our previous paper (11). After removing duplicates, the list consisted of 138 proteins. Qlucore Omics Explorer (Qlucore Omics Explorer Version 3.9) was used for multigroup (ANOVA) comparisons displayed as hierarchical clustering heat maps. For HMC-1 the p-value was set at 0.001, generating a q-value of 0.0020611, and then contained 84 proteins. For THP-1 the p-value was also set at 0.001, generating a q-value of 0.0013661, and then contained 124 proteins.

### 2.6. Lipidomics

Lipid extraction and analysis were performed at the Department of School of Public Health and Community Medicine at Sahlgrenska Academy, University of Gothenburg, Sweden. The lipids analysed and how they were identified are listed in **Supplementary Table 1**.

For the lipidomic analysis, four independent biological replicates were used for each of the four sample types, resulting in 16 samples per cell line and 32 samples in total (n=4 for each vesicle type and cell line). Additionally, a media-only control was created. Briefly, this was achieved by using media supplemented with 10% EV-depleted FBS. This complete media, which had not been in contact with cells, was processed through the EVs isolation protocol described in sections 2.2 and 2.3. Hence, we ended up with 4 media-only controls, one for each subpopulation of EVs.

#### 2.6.1. Lipid extraction

A Bligh and Dyer liquid extraction procedure was used to extract the lipids from the EV preparations. Glass tubes (CORNING 99449-13) were prepared with 1 mL Dichloromethane (DCM) (EMSURE®; Merck), 2 mL Methanol (LC-MS grade LiChrosolv®; Merck) and 790 µL of 0.9wt% NaCl(aq). To the tubes 10 µl of vesicle sample was added, followed by addition of internal standards. The internal standard SPLASH LipidoMIX™ (SKU: 330707, Avanti Polar Lipids, Inc. USA) was used as internal standard for the lipid classes PC/LPC/PE/LPE/SM/PG/PI. The isotope labeled standard GlcCer(d18:1(d5)/18:0) (SKU: 860638, Avanti Polar Lipids, Inc. USA) was used as internal standard for the Ceramide and Hexosylceramide lipid classes. The SPLASH stock standard was diluted 200 times in methanol and 20 µl was added to each sample. The GlcCer(d18:1(d5)/18:0) standard was diluted to 0.027 µg/ml in methanol before adding 10 µl to the sample tubes. The tubes were vortexed for 30 s, ultrasonicated for 5 minutes and put on a shaker for 15 minutes at RT to solubilise the lipids in the monophase. For phase separation 1 mL DCM and 1 mL 0.9wt% NaCl(aq) was added to the extraction tubes, final ratio NaCl(aq):MeOH:DCM [1.8:2:2]. The tubes were vortexed for 30 s and centrifuged at 300 × g for 10 minutes at 10°C to assist the phase separation into two layers. From the lower organic phase 1000 µL was transferred to an Eppendorf tube using a Hamilton syringe (washed 2x in isopropanol and 2x in methanol between samples). The Eppendorf tube was centrifuged at 10 000 × g for 10 minutes at 10°C to remove particulates. From the Eppendorf tubes, 800 µl of each sample was transferred to a HPLC vial (Waters Total Recovery SKU: 186002805) using a CTC-PAL robot configured with a Hamilton syringe. Then solvents were evaporated under a stream of nitrogen gas with a temperature of 30°C (Porvair MiniVap Gemini) in the total recovery vial. The remaining lipid film was solvated with 300 µL acetonitrile/isopropanol (ratio [2:1]). The tubes were vortexed for 30 s, ultrasonicated for 5 minutes, and put on a shaker for 15 minutes at RT to solubilise the lipids fully in the injection solvent.

#### 2.6.2. UHPLC-ESI-QqQ-MS/MS analysis

An initial pre-run was done to determine the optimal dilution for each sample. This was a necessary step due to the large differences in the initial concentrations between samples. The additional dilution step was only necessary for the PC and SM lipid classes. Once the optimal dilutions were determined the study data was collected using this dilution.

Analysis was performed using Ultra High-Performance Liquid Chromatography (Waters ACQUITY UPLC I-Class PLUS) using Electro Spray Ionization and a Triple Quadrupole mass detector (UHPLC-ESI-QqQ-MS/MS) (Waters Xevo TQXS). The mass spectrometer was operating in MRM mode (multiple reaction monitoring) for data acquisition.

Chromatography was performed based on two different separation principles: Hydrophilic interaction chromatography (HILIC) (BEH Amide VanGuard FIT Column, SKU:186009508) and Reverse phase chromatography (RP) (Premier BEH C8 VanGuard FIT Column, SKU:186010360). HILIC method was used for lipids in the classes LPC/PG/PE/SM/PI in order the have chromatographic lipid class separation. PC lipids were analysed on RP due to the large number of species in this class it was desirable to separate them based on the fatty acyl chains.

The lipids Cer and HexCer lipids were analysed on the same column and mobile phase as the PC lipids but with slightly extended gradient. For details on the chromatography mobile phases and gradients see supplementary data.

Lipid panel for targeted UHPLC-ESI-QqQ-MS/MS method were selected based on previous screening experiments. Targeted screening was conducted using Waters LipidQuan MRM database containing over 2000 lipids. To verify that no major lipid was missed additional screening was done using precursor ion scanning for the different lipid class specific fragments (PIS184, PIS264, NL141, NL189, NL277).

#### 2.6.3. Data analysis and quantification

Masslynx 4.2 software was used for data acquisition and Targetlynx was used for peak integration. Integrated peaks were exported to the SAS 9.4 software for quantification. The online tool LICAR was used for isotope correction of co-eluting peaks in HILIC (40). The calculated type II isotope correction factors were applied to the results to account for the isotopic overlap of lipids that are close in mass and have similar retention time. Quantification was made relatively to the internal standard of the same class. Both the Ceramide analytes and the Hexosylceramides were quantified using the GlcCer(d18:1(d5)/18:0) internal standard as they have showed to have very consistent and comparable recoveries. Results were normalized as mol%, meaning that all lipids in a sample were summed up and subsequently expressed as the percentage of the total.

#### 2.6.4. Statistics and bioinformatics

Qlucore Omics Explorer (Omics Explorer Version 3.9) was used for principal component analysis and multigroup comparisons displayed as hierarchical clustering heat maps. GraphPad Prism 10 was used to display differential enrichment of lipid classes.

### 2.7. Nano-flow cytometry

The presence of tetraspanins (CD63, CD81, and CD9) and ADAM10 on the surface of EVs was determined utilizing a Flow NanoAnalyzer (NanoFCM Inc.) following the manufacturer’s guidelines and using a staining protocol as described previously (15, 16). For experimental design, the MIFlowCyt-EV guidelines were consulted (17, 18). Controls, including buffer-only controls, buffer with reagent controls, unstained sample controls, single-stained sample controls, and detergent treatment, were performed. For the detergent treatment control, stained samples, diluted 10-fold, were treated with Triton X-100 and compared to the non-treated sample, and the ratio of disrupted particles was calculated. In this project, we used 0.1% TritonX-100 and incubation for 5 minutes at RT (19).

#### 2.7.1. Instrument setup

The system was calibrated before samples were loaded and analyzed. For alignment, a 1:100 dilution of 0.25 μm Fluorescent Silica Microspheres (250 nm Std FL SiNPs) was used at a laser power of 20/50 mW 488 & 20/100 mW 638 at a side scatter (SS) decay of 0.2%. The 250 nm Std FL SiNPs bead data was recorded and analysed as the concentration standard using the large signal auto threshold. S16M-Exo Silica Nanosphere Cocktails were used to create a calibration curve of particle sizes and side scatter intensity. S16M-Exo includes particles at a range of 68 - 155 nm (peaks at 68, 91, 113, 155 nm) and were recorded at a laser power of 10/50 mW 488 & 20/100 mW 638; SS decay of 10% and analysed using the small signal auto threshold. The pressure was kept at 1.0kPa for all standards and samples.

#### 2.7.2. Staining

First, the concentration of the EV samples was determined to ensure that the stained samples would have the recommended concentration of 2,000 and 12,000 events per minute. Based on these measurements, EV samples were pre-diluted before antibody staining. Prediluted antibodies (**Table 4**), isotype controls (**Table 5**), or PBS were added to 3 µl pre-diluted EV samples to a final volume of 5 µL. The samples were incubated in the dark for 40 minutes. After incubation, 45 µL PBS (0.0067 M PO_4_, pH 7.0 to 7.2, Cytiva, HyClone Laboratories) was added to the samples, leading to a 10X dilution of the EV-antibody mix. Directly before acquisition, the samples were further serial diluted to reach final dilutions of 100x and 200x before recording on the nFCM instrument. Dilutions were done with TE Buffer (Tris-EDTA, 1X Solution, pH 7.4, Molecular Biology, Fisher BioReagents, US).

**Table 4.**
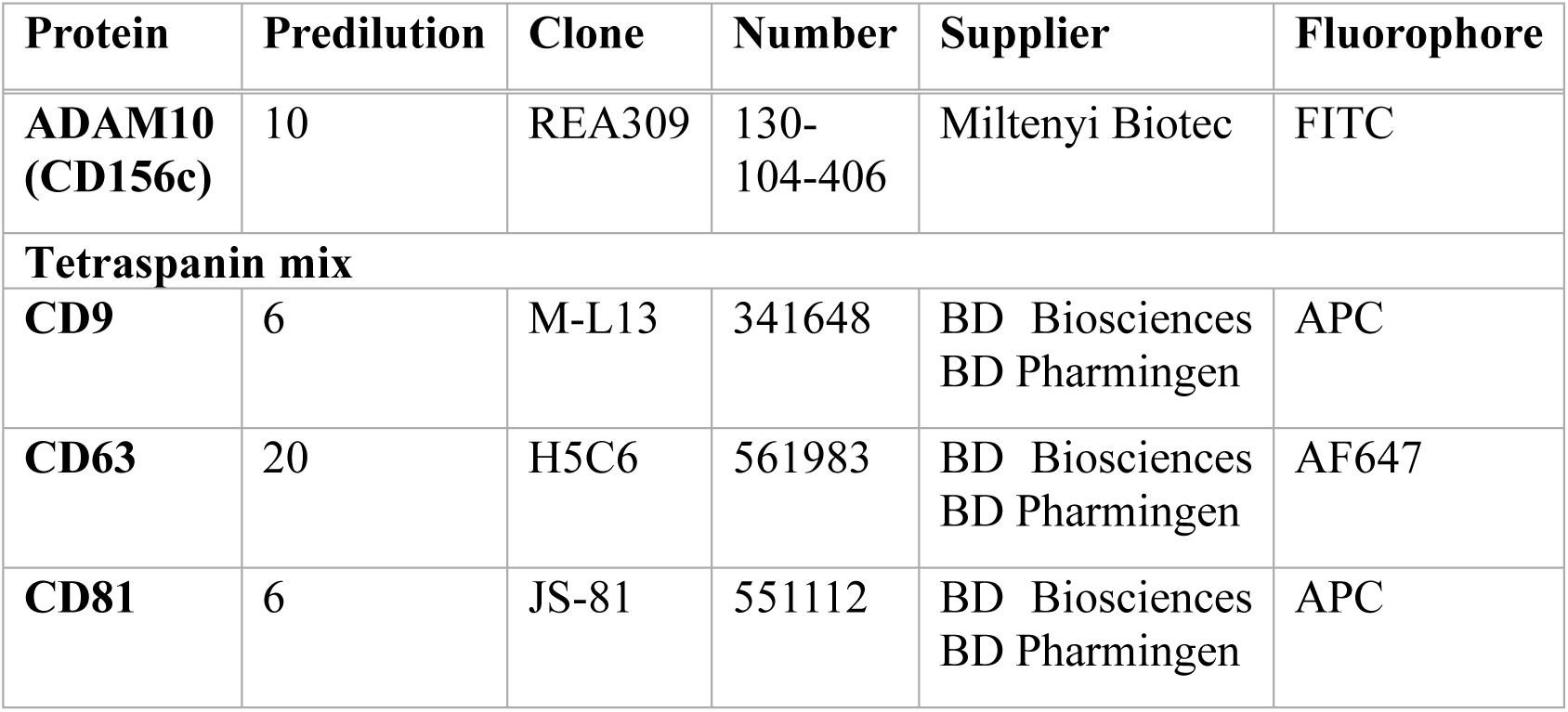
Antibodies and their predilutions used for Flow NanoAnalyzer.

**Table 5.** Isotype control antibodies and their predilutions used for nFCM.

| **AB** | **Isotype control for:** | **Predilution** | **Clone** | **Number** | **Supplier** | **Fluorophore** |
| --- | --- | --- | --- | --- | --- | --- |
| **REA Control Antibody (S), human IgG1, FITC** | ADAM 10-FITC | 33 | REA293 | 130-113-437 | Miltenyi Biotec | FITC |
| **APC Mouse IgG1, κ Isotype Control** | CD9-APC and CD81-APC | 6 | MOPC-21 | 555751 | BD Biosciences<br>BD Pharmingen | APC |
| **AF647 Mouse IgG1, κ Isotype Control** | CD63-AF647 | 20 | MOPC-21 | 557714 | BD Biosciences<br>BD Pharmingen | AF647 |

#### 2.7.4. Acquisition

Samples were run at the same sampling pressure as the concentration standard and with the same laser power as the size standards. The sampling pressure was set to 1.0 kPa. The side scatter was set as the triggering channel (each signal above the SSC threshold was acquired), and samples were analysed using the auto threshold according to the EXO settings.

#### 2.7.5. Analysis, statistics and bioinformatics

The nano-FCM software (NF profession V1.0) was used to calculate the percentage of positive signal, particle concentration, and size distribution. To calculate the percentage of positive events, particle concentration, and size distribution the NanoFCM software (NanoFCM Profession V2.0; NanoFCM, Inc.) was used. The auto threshold (small) was set for each sample, and gates were set.

### 2.8. Data availability

The MS proteomics data have been deposited to the ProteomeXchange Consortium via the PRIDE partner repository with the dataset identifier PXD063334 (20, 21). We have submitted all relevant data from our experiments to the EV-TRACK knowledgebase (EV-TRACK ID:EV250057 (22).

## 3. Results

### 3.1. Immune cells secrete EVs of different sizes and densities

We have previously determined the protein cargo of two EV subpopulations of different sizes released by breast cancer cells (1). In the current study, we build on these findings by determining the protein and lipid content in four immune cell-derived EV subpopulations. For this purpose, we expanded on the isolation method by incorporating a density cushion that enables the isolation of EV subpopulations according to density. This resulted in the determination of the proteomic and lipidomic profile of four subpopulations of EVs enriched at different sizes and densities.

Conditioned medium was collected from the immune cell lines HMC-1 and THP-1, and crude L-EVs and S-EVs were isolated by differential ultracentrifugation (**Figure 1A**). Subsequently, a bottom-loaded density cushion centrifugation was used to purify and sub-fractionate the crude samples further. As EVs were collected at two different densities for each crude EV type (**Supplementary** Figure 1), this resulted in four types of EVs: L-EVs low density (L-EVs LD), L-EVs high density (L-EVs HD), S-EVs low density (S-EVs LD), and S-EVs high density (S-EVs HD) (**Figure 1A**). The LD EVs were collected in the interphase between 10 and 24% Iodixanol (1.079-1.146 g/ml), and HD EVs were collected in the interphase between 24 and 32% Iodixanol (1.146-1.185 g/ml). NTA and protein measurements showed that the yield was generally the highest for L-EVs LD, and the lowest for L-EVs HD (**Supplementary** Figure 2A-B). Furthermore, TEM showed that all four EV subtypes indeed contained EVs (**Figure 1B-C** and **Supplementary** Figure 3). To be able to accurately determine the different EV sizes in the four subpopulations, we conducted a double-blinded experiment where several thousand EVs were manually annotated (**Figure 1D-E**). L-EV fractions generally contained a higher amount of bigger EVs than S-EV fractions. Additionally, HD fractions contained EVs that were smaller than the corresponding LD fraction. Notably, L-EVs LD were more heterogeneous in size, and these samples also had fewer non-EV particles than L-EVs HD. Overall, S-EVs were smaller than L-EVs, additionally, more non-EV particles could be seen in these samples. The most notable difference between S-EVs LD and S-EVs HD was the high amount of non-EV particles in the HD factions.

To characterise the four EV subtypes, we evaluated the presence of a general EV marker (flotillin-1), a non-EV protein (calnexin), and markers that have been suggested to be enriched in either S-EVs (CD63, CD9, and CD81) or L-EVs (mitofillin and mitochondrial import receptor subunit TOM20 homolog [TOMM20]) with western blot. All four EV subtypes were positive for flotillin-1 (**Figure 1F**), which confirms our previous findings in breast cancer-derived EVs that all EV subtypes contain flotillin-1 to a similar degree (1). As expected mitofillin and TOMM20 showed L-EV enrichment (**Figure 1G**). The tetraspanins CD9, CD63, and CD81 were present in all fractions, however, an S-EVs enrichment could be observed, which was more prominent in the HMC-1 S-EVs (**Figure 1H**). A signal for calnexin was mainly detected in THP-1 L-EVs (**Figure 1I**). These results, together with the TEM results, suggest that we have enriched EVs with different sizes, densities, and protein cargo.

### 3.2. The proteomes of the four EV subpopulations are significantly different

We performed quantitative proteomics to identify differentially enriched proteins in the four subpopulations of EVs, which were all analysed in biological triplicates for each cell line. In total, 5364 proteins were quantified in our dataset, 3970 and 4736 of these proteins were quantified in HMC-1 and THP-1 cell line-derived EVs, respectively, with 3342 being quantified in both. To visualize the relationship between the different types of isolated EVs, a principal component analysis (PCA) was performed for each cell line, including all proteins. Component 1, representing around 50% of the variability, distinguished L-EVs and S-EVs (**Figure 2A-B**). Furthermore, components 2 and 3 distinguished the LD and HD fractions; however, this was more evident in the THP-1-derived EVs as compared to the HMC-1-derived EVs. A similar pattern was observed when all samples from both cell lines were analysed in the same PCA plot, although the samples were then also separated based on the cell type in components 2 and 3 (**Figure 2C**).

**Figure 2.**
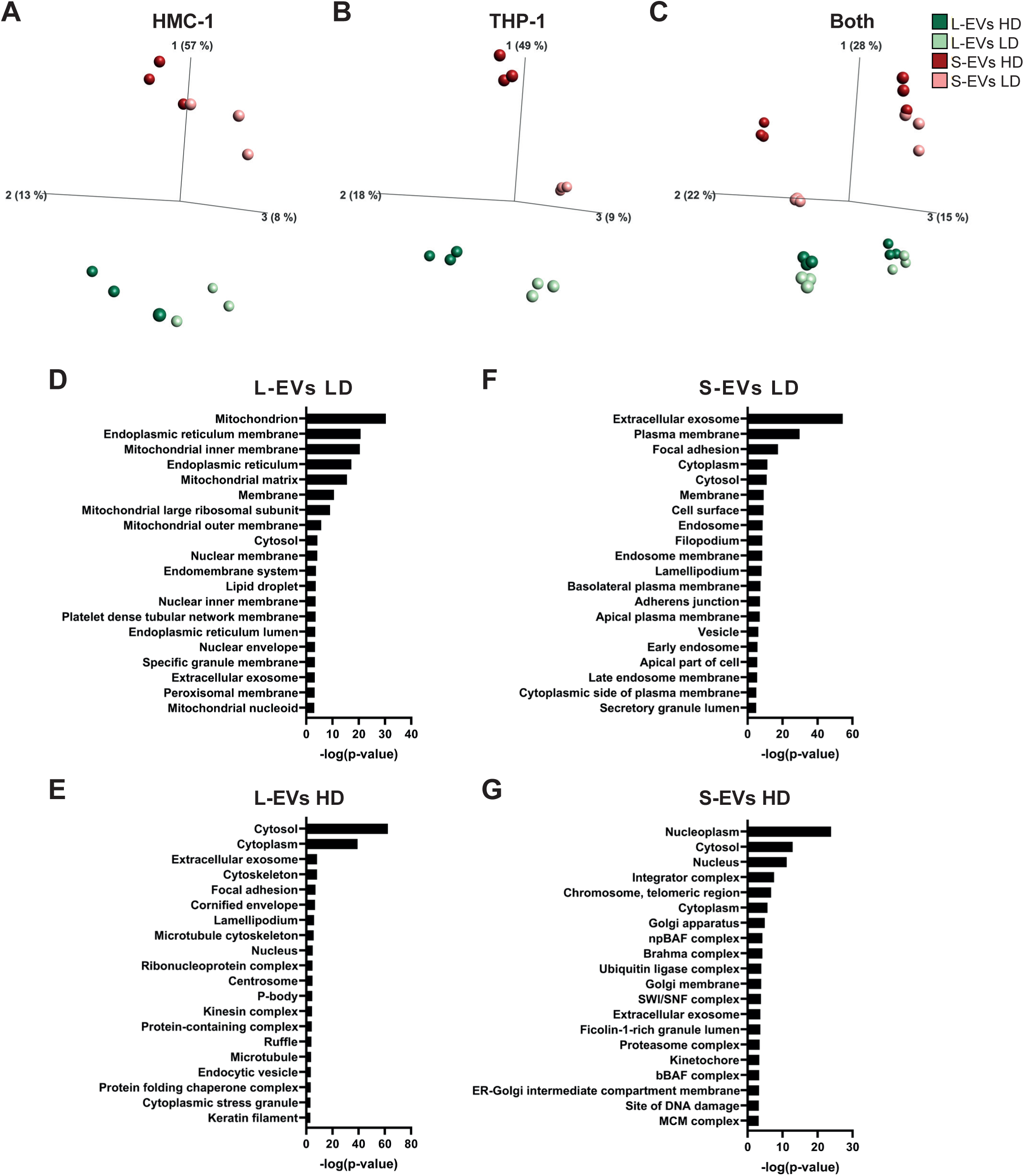
The proteome of the four EV subpopulations is substantially dissimilar. Quantitative proteomics (tandem mass tag) was used to determine the differences in the proteomes of the EV subpopulations. Three biological replicates (40μg protein/sample) were used from both cell lines. **(A-C)** Principal component analysis illustrating the relationship between the subpopulations derived from HMC-1 (A), THP-1 (B) and both cell lines combined (C). **(D-G)** Database for Annotation, Visualization and Integrated Discovery (DAVID) was used to determine the most enriched cellular compartments associated with proteins enriched in the four different EV subpopulations. The 15 most enriched terms (based on p-value) for each EV subtype are displayed. The enriched proteins were produced as described in **Supplementary** Figures 4 and 5. HD, high density; LD, low density; L-EVs, large extracellular vesicles; S-EVs small extracellular vesicles

Next, we performed pairwise comparisons between the different EV subtypes, revealing that proteomic differences among EV subpopulations were more pronounced by size than by density (**Supplementary** Figure 4A-J). Proteins identified as uniquely upregulated in EV subpopulations (**Supplementary** Figure 5A-D) were analysed with the database for annotation, visualization, and integrated discovery (DAVID) to determine the most enriched biological cellular compartments associated with these proteins (**Figure 2D-G**). Proteins upregulated in L-EVs LD were associated with mitochondrion, endoplasmic reticulum, and membrane, while the proteins upregulated in L-EVs HD were associated with cytosol, cytoskeleton, kinesin complex, and microtubule (**Figure 2D-E**). The proteins upregulated in S-EVs LD were associated with the cytoplasm, plasma membrane, and early and late endosome, while the proteins upregulated in S-EVs HD were associated with the nucleus, cytosol, and Golgi apparatus (**Figure 2F-G**). The GO term “extracellular exosome” was associated with the upregulated proteins for all subpopulations, but had the highest enrichment in S-EVs LD.

Taken together, L-EVs and S-EVs were well separated, however, the separation between the LD and HD samples was more pronounced for the THP-1 EVs than for the HMC-1 EVs. In addition, the differently enriched proteins in the EV types were associated with different biological functions and organelles. Both the PCA plots and the GO term enrichment analysis strengthen the assumption that our four sample types are enriched for different subpopulations of EVs. This indicates that our EV subtypes might have distinct biogenesis and biological functions, which could aid in elucidating the differences between EV subpopulations.

### 3.3. Immune cells secrete EVs with different expression of common EV markers

The International Society for Extracellular Vesicles (ISEV) have published three Minimal Information for Studies of Extracellular Vesicles (MISEV) guidelines, MISEV2014, MISEV2018, and MISEV2023 (13, 23, 24). In the MISEV2018 and MISEV2023, five categories of proteins were presented to be used when analysing the protein content of EVs. Categories 1 and 2 asses the presence of EVs, with Category 4 providing additional information on intracellular origins. Categories 3 and 5 asses the presence of contaminations and co-isolates. We analysed all the proteins quantified in our dataset across these five categories. Category 1 assesses the presence of EV membrane proteins from the plasma membrane (PM) and the endosomal pathway. The majority of the tetraspanins, ADAM10, and some integrins were enriched in S-EVs LD in THP-1 and in S-EVs in HMC-1, while syndecans and glypicans were enriched in both S-EVs LD and S-EVs HD in both cell lines (**Figure 3A** and **Supplementary** Figure 6A). Furthermore, major histocompatibility class I and II, heteromeric G proteins, and the rest of the integrins were enriched in both L-EVs LD and S-EVs LD (**Figure 3A** and **Supplementary** Figure 6A). Belonging to Category 2 (cytosolic EV-associated proteins), the majority of the ESCRT proteins were enriched in S-EVs LD, while SDCBP (syntenin-1) was enriched in both S-EVs LD and S-EVs HD (**Figure 3B** and **Supplementary** Figure 6B). Flotillin-1 and −2 were enriched in both L-EVs LD and S-EV LD, while the tubulins were either enriched in L-EVs HD or S-EVs HD (**Figure 3B** and **Supplementary** Figure 6B). In Category 3 proteins of non-EV co-isolated structures are summarized. The 14-3-3 proteins (YWHAs) were enriched in S-EV LD and TGFB1 was enriched in L-EVs LD, while the rest of the proteins were enriched in S-EVs HD or in both S-EV subtypes (**Figure 3C** and **Supplementary** Figure 6C). In Category 4 the proteins associated with mitochondria, ER, and Golgi apparatus were enriched in L-EVs LD, while the nucleus proteins such as histones were enriched in S-EV HD (**Figure 3D** and **Supplementary** Figure 6D). Lastly in Category 5, which contains “secreted proteins recovered with EVs”, the majority of proteins were enriched in S-EV HD, except for IL-18 which was enriched in L-EV LD and L-EV HD (**Figure 3E** and **Supplementary** Figure 6E). Together these results demonstrate that the EV-associated proteins (categories 1 and 2) are mainly associated with S-EV LD and to some extent L-EVs LD, the proteins more associated with contamination/co-isolates (categories 3 and 5) are enriched in S-EV HD. Furthermore, the proteins associated with organelles other than the plasma membrane and endosomes (category 4) are enriched in L-EVs LD, except for histones, which were enriched in S-EVs HD.

**Figure 3.**
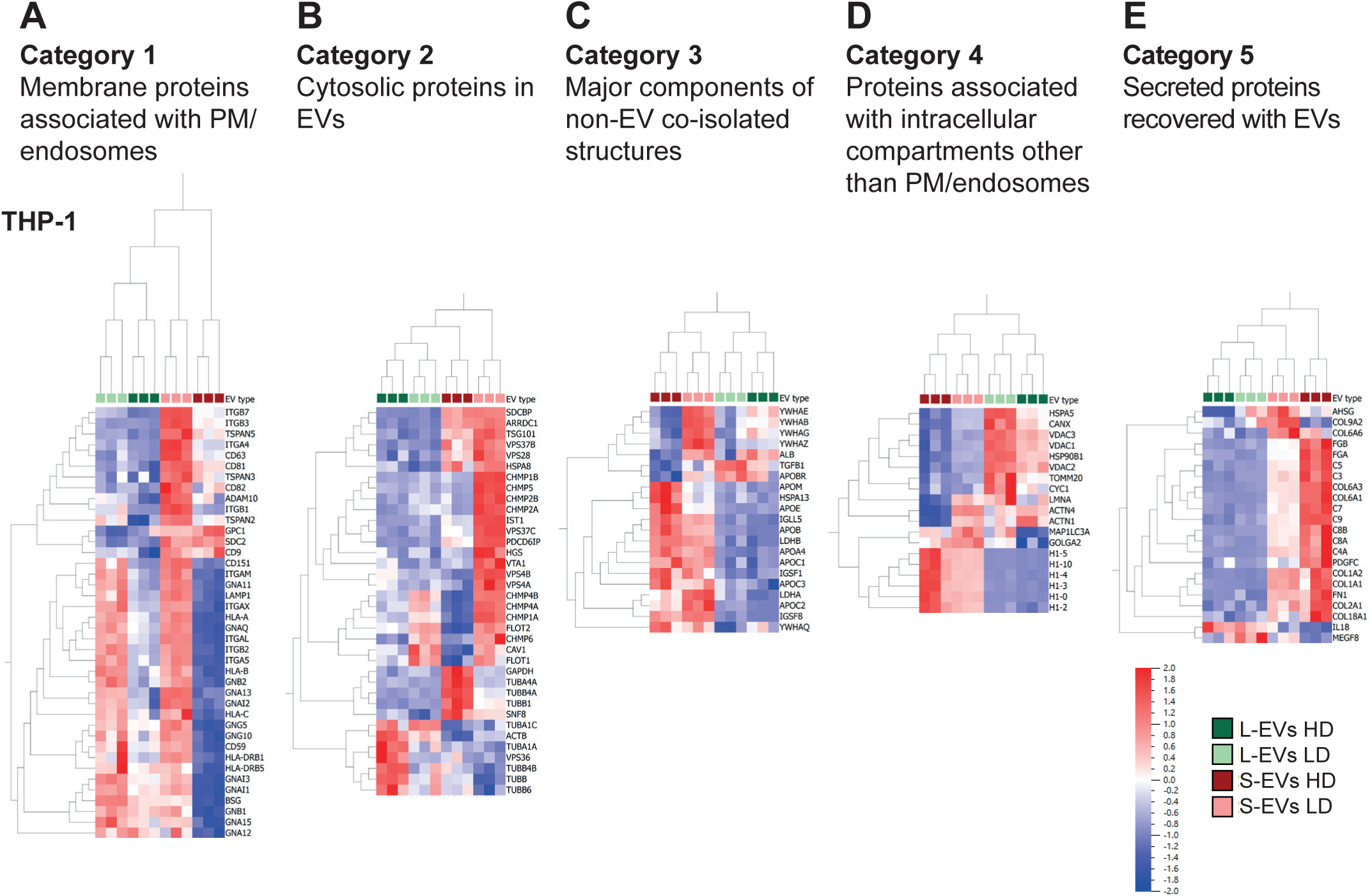
The expression of MISEV guideline proteins in the four EV subpopulations from THP-1 cells. **(A-E)** In the MISEV guidelines (13, 24), five categories of proteins are recommended to be evaluated in EV isolates. We have analysed all proteins in our dataset that belong to these five categories. (**A**) Category 1 – Membrane proteins associated with the plasma membrane (PM)/endosomes. (**B**) Category 2 – Cytosolic proteins in EVs. (**C**) Category 3 – Major components of non-EV co-isolated structures. (**D**) Category 4 – Proteins associated with intracellular compartments other than plasma membrane (PM)/endosomes. (**E**) Category 5 – Secreted proteins recovered with EVs.

To further determine the expression of common EV proteins in our data set, we next constructed a list of previously identified markers for large and small EVs, discovered by us and others (see method section) (**Figure 4A-B**). This analysis showed that proteins such as ADAM10, CD63, CD81, PDCD6IP (ALIX), TSG101, and 14-3-3 proteins (YWHAs) and EZR (Ezrin) were enriched in S-EV LD in THP-1 (**Figure 4B**, cluster 1), while proteins such as GAPDH, APOE, and Chaperonin Containing TCP-1 complex proteins (CCTs) were enriched in S-EVs HD in THP-1 (**Figure 4B**, cluster 2). For HMC-1, these proteins were enriched in both S-EV subpopulations (**Figure 4A**, cluster 1). SDCBP (Syntenin-1) was enriched in both S-EVs LD and HD in both cell lines (**Figure 4A**, cluster 1, **Figure 4B**, cluster 3). Proteins such as TOMM22, MICOS complex subunits (IMMT [mitofilin], CHCHD3), heat shock proteins (HSP*), ATP synthase subunits (ATP5s) were enriched in L-EVs LD in THP-1 (**Figure 4B**, cluster 4). Some of these proteins were also enriched in L-EVs LD in HMC-1, while others were enriched in both L-EVs LD and L-EVs HD (**Figure 4A**, clusters 2 and 3). Lastly, kinesins (KIFs), PRC1, CIT, RACGAP1, and tubulins (TUBs) were enriched in L-EVs HD for both cell lines (**Figure 4A**, cluster 4, **Figure 4B**, cluster 7). Annexins (ANXAs), Rab proteins (RABs), and flotillin-2 were enriched in both S-EV LD and L-EV LD in THP-1 (**Figure 4B**, cluster 6), while they were not significantly different in HMC-1 suggesting that these proteins are equally expressed in L-EVs and S-EVs. This shows that we could further separate some of the markers that have previously been suggested to be enriched in L-EVs or S-EVs to also be enriched in EVs at certain densities. Together these data demonstrate that L-EVs LD are enriched in proteins associated with the mitochondrion such as TIMMs/TOMMs, MICOS complex subunits, and ATP synthase subunits, while L-EVs HD are enriched in KIFs and proteins associated with cytokinesis and microtubule (**Figure 2-4** and summarized in **Figure 5**). Furthermore, S-EVs LD were enriched in proteins associated with the plasma membrane and the endosomal pathway, such as tetraspanins, ADAM10, proteins from the ESCRT machinery, and 14-3-3s, while S-EVs HD were enriched in tubulins, histones, complement factors, and Chaperonin Containing TCP-1 complex proteins. Additionally, some proteins were enriched in two or more EV subtypes, such as flotillin-1 which was similarly expressed in all EV subtypes, syntenin-1 being enriched in both S-EV subtypes, and RABs, annexins, some integrins, and heteromeric G proteins that were enriched in both LD subtypes (**Figure 5**).

**Figure 4.**
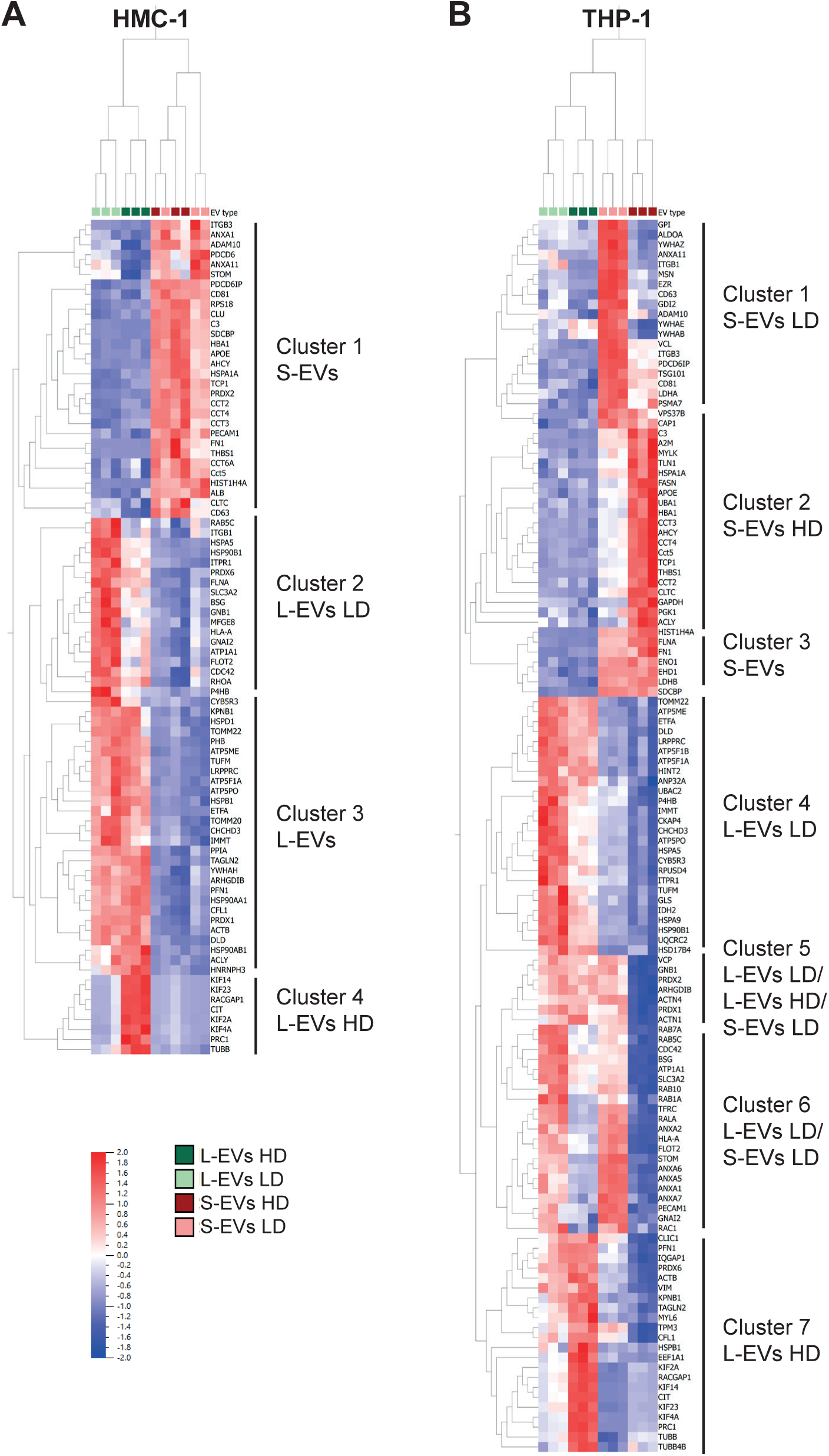
Presence of classical EV markers in EV subpopulations from immune cells. The top 100 proteins identified in EVs from two online EV databases – EVpedia and VesiclePedia – were downloaded (73, 74). Additionally, markers suggested by Lim and colleagues were added to the list (14). Lastly, we also added proteins enriched in L-EVs in our previous paper (11). After duplicates were removed, the list contained 181 proteins. **(A-B)** A multi-group (Anova) comparison was performed in Qlucore and showed that 84 and 124 of these proteins were differentially expressed in EVs from HMC-1 (p-value = 0.001, q-value = 0.0020611) (A) and THP-1 (p-value = 0.001, q-value = 0.0013661) (B), respectively.

**Figure 5.**
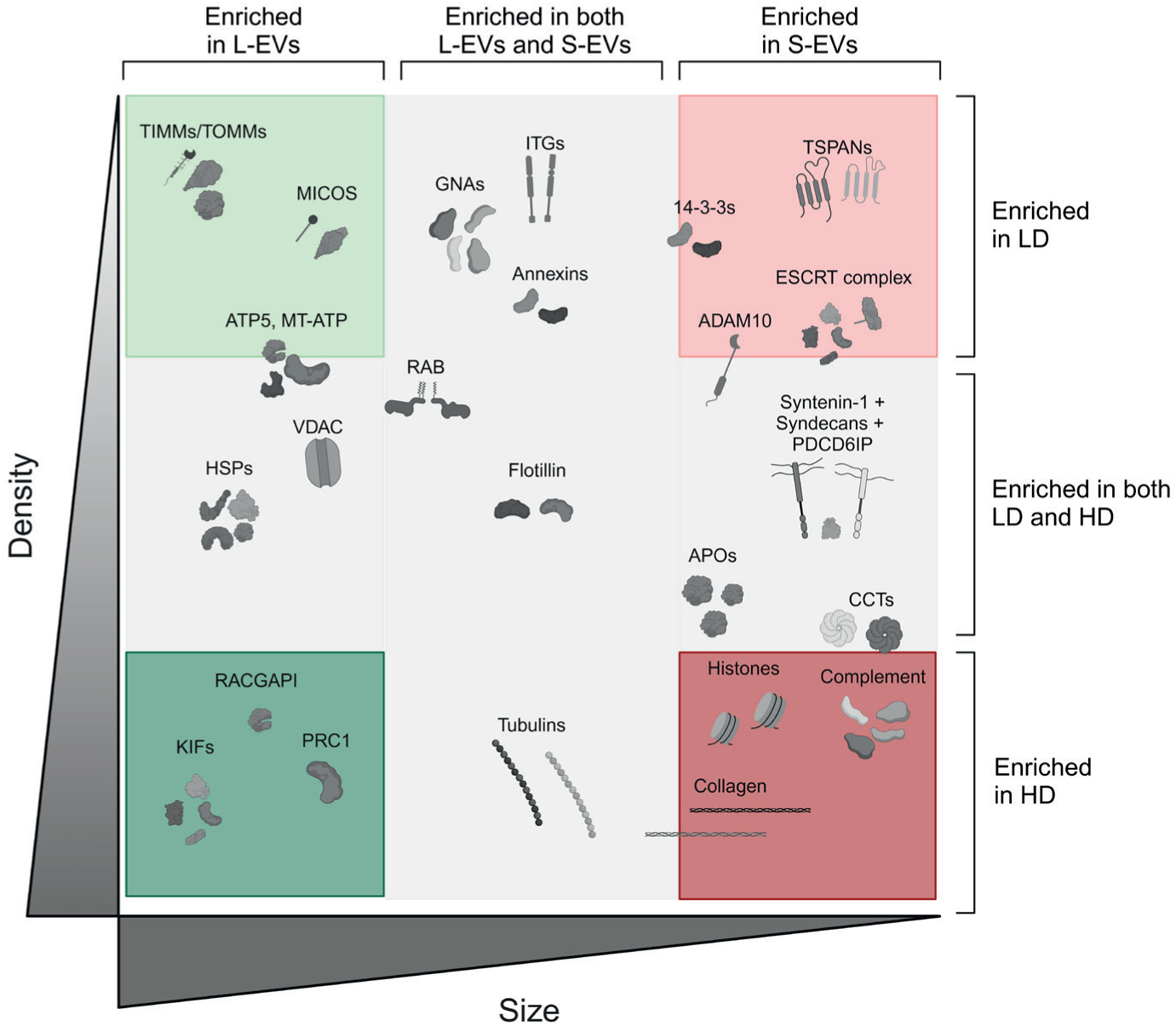
Proteins enriched in subpopulations of EVs. Schematic illustration showing the proteins enriched in the four subpopulations of EVs as determined by quantitative proteomics. Proteins enriched in L-EVs LD, L-EV HD, S-EVs LD, and S-EVs HD are shown in light green, dark green, light red, and dark red boxes, respectively. Proteins in the grey area are enriched in two or more subpopulations. APO, apolipoproteins; CCT, chaperonin containing TCP-1 proteins; ESCRT, endosomal sorting complexes required for transport; GNAs, Guanine nucleotide-binding proteins; HD, high density; HSP, heat shock proteins; ITG, integrins; KIF, kinesin-like proteins; LD, low density; L-EVs, large extracellular vesicles; MICOS, mitochondrial contact site and cristae organising system; RAB, RAB G-proteins; S-EVs small extracellular vesicles; TIMM, translocase of the inner membrane of the mitochondria; TOMM, translocase of the outer membrane of the mitochondria; TSPANs, tetraspanins; VDAC, voltage-dependent anion channels.

### 3.4. ADAM10 is expressed on tetraspanins^+^ EVs

As we observed that CD9, CD63, CD81, and ADAM10 were all enriched in S-EVs (**Figure 3A**, **Figure 4A-B**), and it has previously been shown that ADAM10 is associated with tetraspanins on the cell membrane and that the tetraspanins can regulate ADAM10 activity and its substrate cleavage (25–27), we decided to analyse all tetraspanins and ADAMs/ADAMTSs in more detail in our dataset. All tetraspanins were enriched in S-EVs with a further enrichment in S-EVs LD in THP-1, except for CD151 which was enriched in L-EV LD for both cell lines (**Figure 6A-B**). For the ADAMs/ADAMTSs, ADAM10 was enriched in S-EVs LD, while ADAM17 was enriched in both S-EV LD and L-EV LD (**Figure 6C-D**). Additionally, in THP-1 the ADAMTSs were mostly enriched in S-EV HD (**Figure 6C-D**). As proteomics is a bulk analysis, we next set out to determine if ADAM10 and tetraspanins are co-expressed on the same EVs. To do so, we analysed S-EVs from the HMC-1 with a flow nanoAnalyzer. As we have observed in previous flow nanoAnalyzer experiments that Iodixanol can interfere with the flow nanoAnalyzer measurements, it was not feasible to analyse the S-EVs LD, instead, we used crude S-EVs to investigate the presence of CD9, CD63, CD81, and ADAM10 on their surface. It was shown that the majority of ADAM10 is expressed on EVs that also express CD9, CD63, and/or CD81 in S-EVs (**Figure 6E**). Importantly, Triton-X treatment disrupted the majority of the ADAM10^+^ and tetraspanins^+^ EVs, suggesting they are true EVs (**Supplementary** Figure 7). All relevant controls for flow nanoAnalyzer can be found in **Supplementary** Figures 7-9.

**Figure 6.**
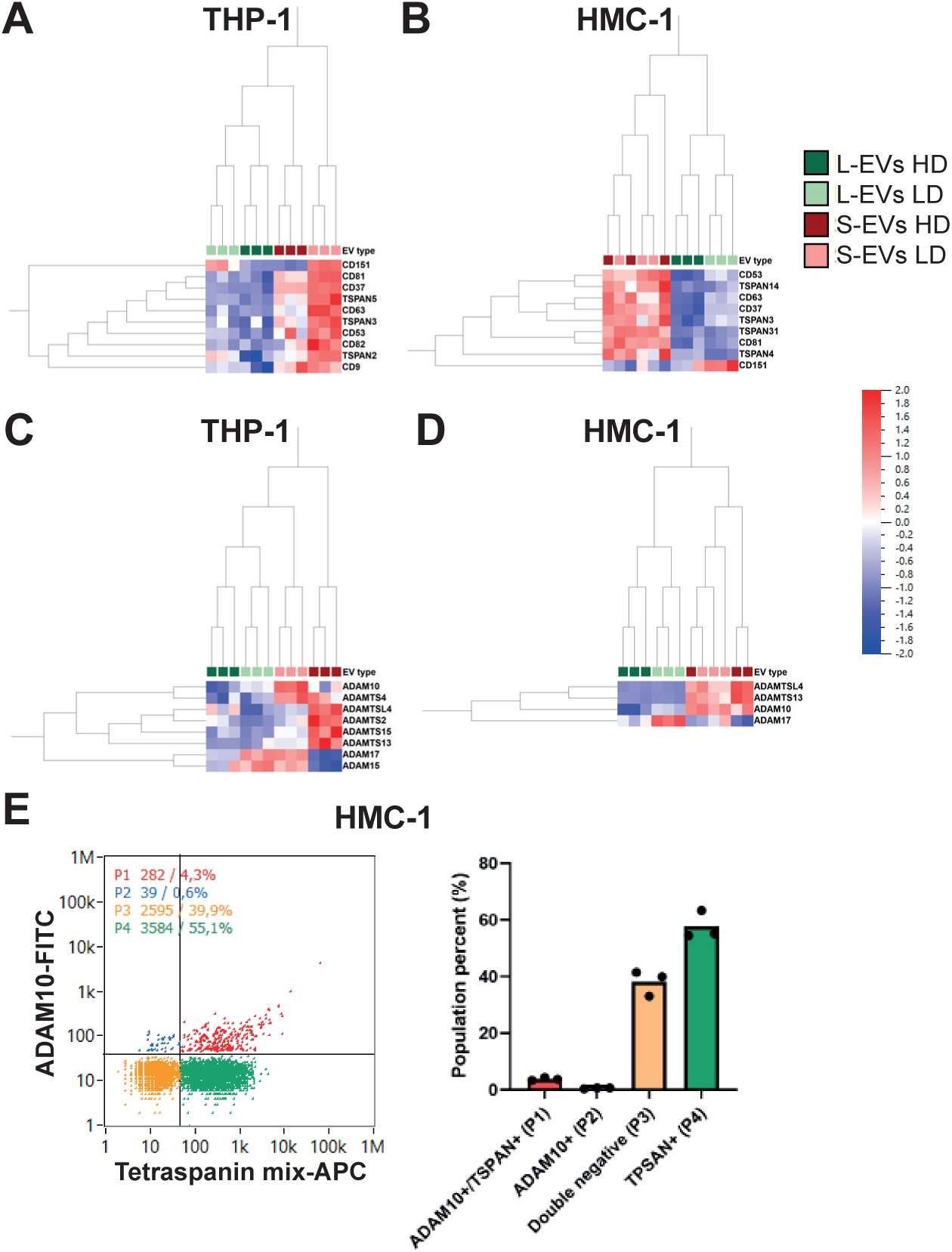
ADAM10 is present on tetraspanins expressing EVs. (**A-D**) A multi-group (Anova) comparison was performed in Qlucore with a focus on the tetraspanins (A-B) and ADAMs and ADAMTSs (C-D). (**E**) NanoFCM was used to determine the co-expression of ADAM10 and tetraspanins on single S-EVs. S-EVs from HMC-1 cells were stained for ADAM10 and an anti-tetraspanins mix containing antibodies for CD63, CD9, and CD81.

### 3.5. The lipidomes of the four EV subpopulations are altered

Next, we sought to determine the lipid composition of the EV subtypes. We performed mass spectrometry to determine the presence of 107 lipids in the four subpopulations of EVs, all analysed in biological quadruplicates for each cell line. Overall, the four media-only blanks contained low levels of lipids. Importantly, they contained substantially less than the corresponding samples, with a median contribution of 0.6% from the media-only blanks to the lipids in the samples. One exception was LPC(18:2_B), which was high in one of the media-only blanks. Interpretation of this lipid should therefore be made with caution.

To visualize the relationship between the different types of isolated EVs, a PCA was performed for each cell line, including all lipids. The samples were well grouped based on EV type and component 1, representing over 60% of the variability in THP-1, distinguished L-EVs and S-EVs (**Figure 7A-B)**. Furthermore, components 2 and 3 distinguished the LD and HD fractions; however, this was more evident in the HMC-1-derived EVs as compared to the THP-1-derived EVs. A similar pattern was observed when all samples from both cell lines were analysed in the same PCA plot, although the samples separated based on cell type (**Figure 7C**).

**Figure 7.**
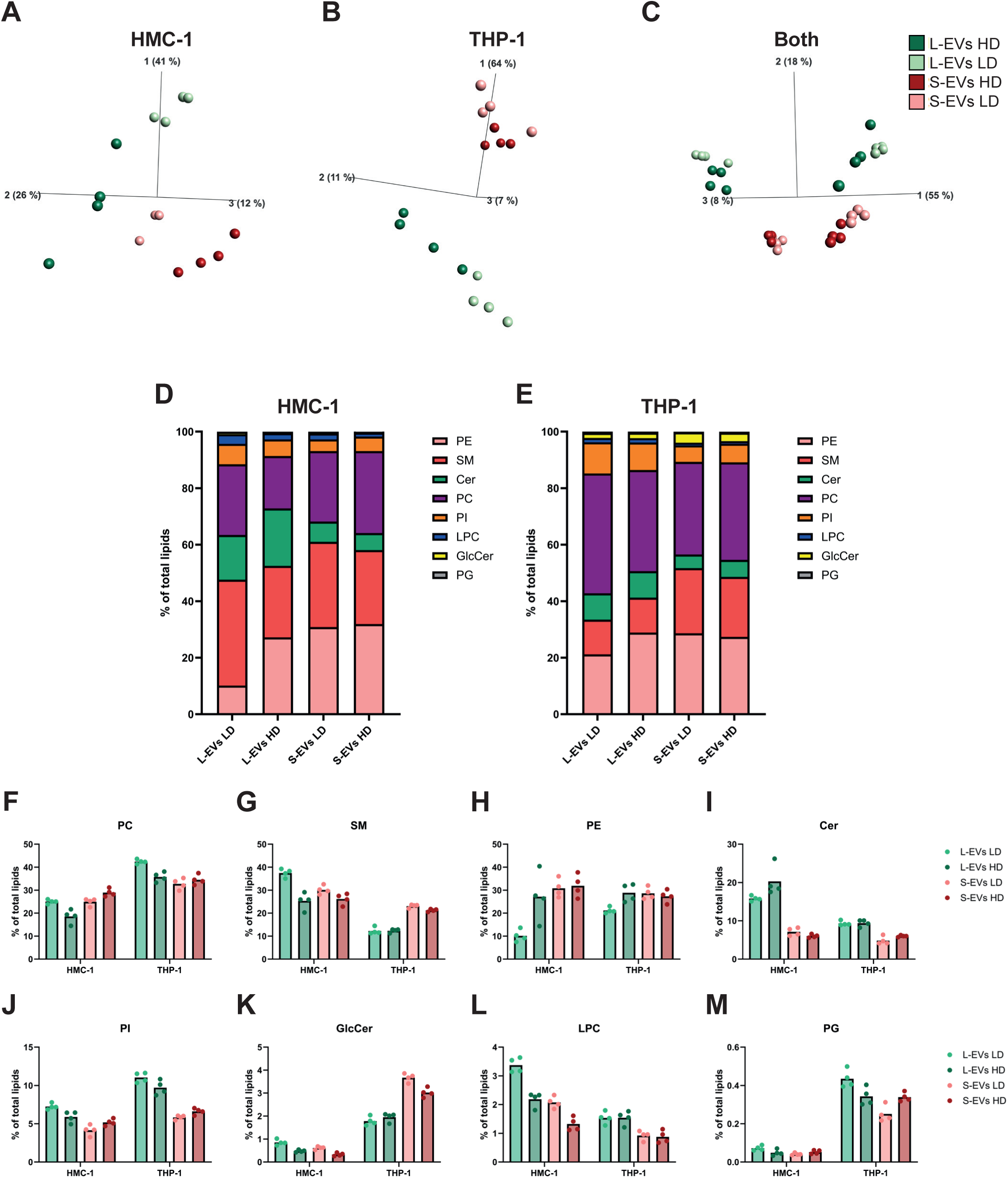
Lipidomic analysis of subpopulation of EVs from immune cells. Mass spectrometry was used to determine the differences between 107 lipids in the EV subpopulations. Four biological replicates (10µl EVs/sample) were used from both cell lines. **(A-C)** Principal component analysis illustrating the relationship between the subpopulations derived from HMC-1 (A), THP-1 (B) and both cell lines combined (C). (**D-M**) The % for each lipid class of the total lipids is illustrated.

Firstly, we analysed the lipids at the group level for each lipid type. Notably, significant differences were observed between the two cell lines. While sphingomyelins (SM) and ceramides (Cer) were more common in HMC-1 EVs as compared to THP-1 EVs, phosphatidylcholine (PC), phosphatidylinositols (PI), glucosylceramide (GlcCer) and phosphatidylglycerol (PG) were more abundant in THP-1 EVs as compared to HMC-1 EVs (**Figure 7D-M)**. Indeed, SM was the most abundant lipid in HMC-1 EVs, whereas PC was the most abundant lipid in THP-1 EVs. Importantly, there were differences between the EV subpopulations; most prominent was the lower abundance of phosphatidylethanolamine (PE) in L-EVs LD compared to all three other EV subpopulations (**Figure 7D-E, H**). In the HMC-1 L-EVs LD this was mainly compensated by an increase in SM, PI, and LPC, while in THP-1 L-EVs LD it was compensated by an increase in PC, PI, and PG (**Figure 7D-M**). Furthermore, Cer, PI, and LPC were more abundant in L-EVs compared to S-EVs, while SM and GlcCer were more enriched in S-EVs at least in the THP-1 EVs (**Figure 7D-M**).

Secondly, we focused on an analysis of individual lipids. We identified the ten most abundant lipids in each EV subtype. In each sample, these ten lipids represented approximately 70% of all the lipids (**Figure 8A**). In the HMC-1 cell line, one of the largest differences was for PE(O-2:0_18:1), a plasmanyl, and PE(O-2:0_18:2), which were less common in L-EV LD. Additionally, Cer(d18:1/22:0) and Cer(d18:1/16:0) were less common in S-EVs compared to L-EVs. In the THP-1 cell line, there was a large difference in PC(16:0/16:0) also called (DPPC), that were much less abundant in L-EVs as compared to S-EVs. Additionally, PE(O-2:0 _18:2) was also less abundant in L-EVs LD, similar to that for HMC-1.

**Figure 8.**
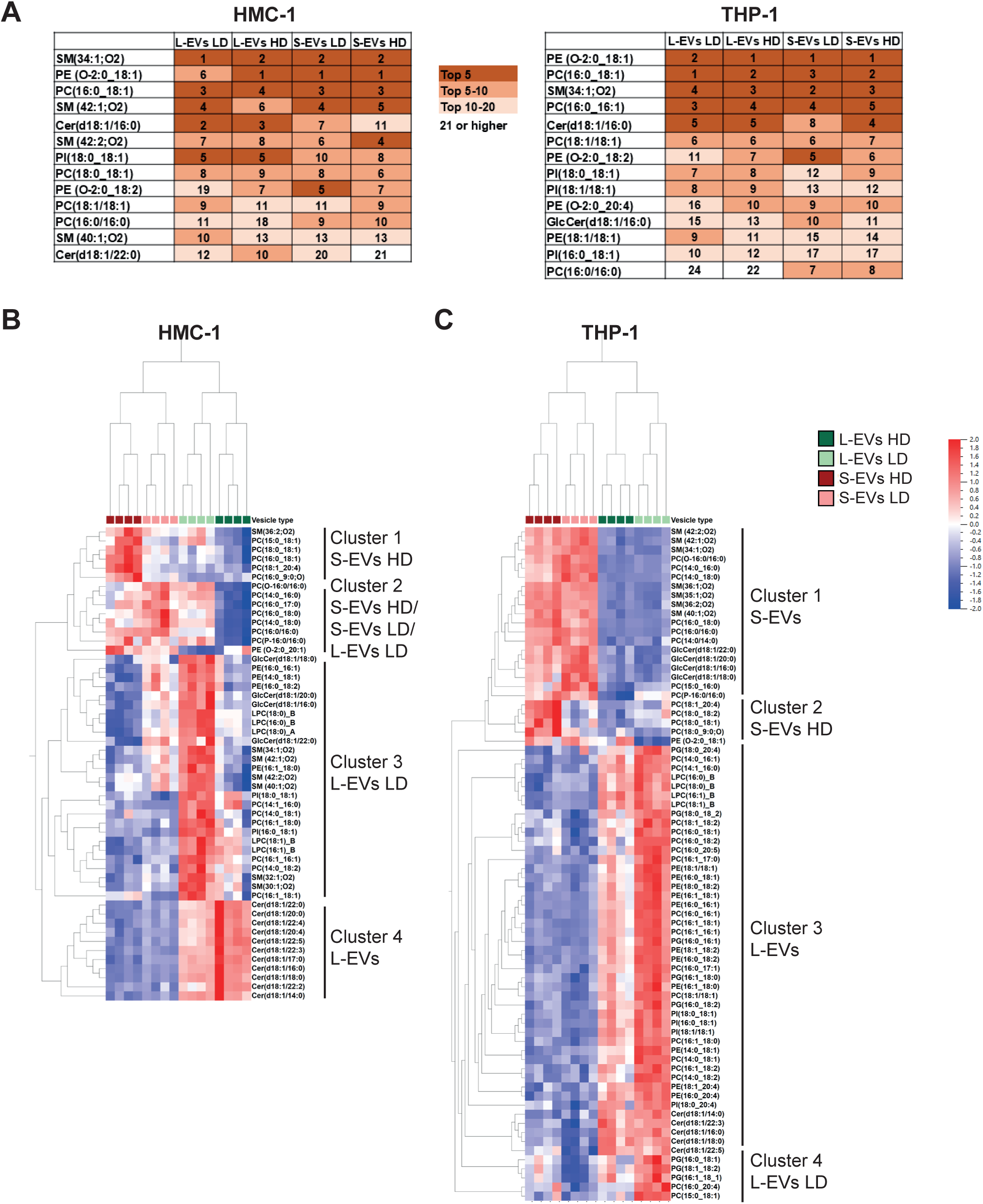
Analysis of individual lipids. (**A**) The top 10 most abundant lipids in each EV subtype and their positions in the other EV subtypes. (**B-C**) A multi-group (Anova) comparison was performed in Qlucore for lipids in EVs from HMC-1 (B) and THP-1 (C).

Focusing on lipids that were commonly altered in both cell lines, analysis of individual lipids also demonstrated that Cer(d18:1/22:5), Cer(d18:1/18:0), Cer(d18:1/16:0), Cer(d18:1/22:3), and Cer(d18:1/14:0) were enriched in both L-EV subpopulations compared to S-EVs (**Figure 8B**, cluster 4 and **8C**, cluster 3). Conversely, SM(36:2;O2) were enriched in S-EVs HD/S-EVs compared to L-EVs (**Figure 8B**, cluster 1 and **8C**, cluster 1) and PC(18:0_18:1) and PC(18:1_20:4) were enriched in S-EVs HD compared to all three other subpopulations (**Figure 8B**, cluster 1 and **8C**, cluster 2). Meanwhile, several of the other PCs were enriched in other subpopulations. Lastly, LPC(18:1)_B, LPC(16:1)_B, PC(16:1_16:1), PC(14:0_18:2), PC(16:1_18:1), PC(14:1_16:0), PC(16:1_18:1), PI(16:0_18:1), and PI(18:0_18:1) was enriched in L-EVs LD/L-EVs (**Figure 8B**, cluster 3 and **8C**, cluster 3).

Together, this suggests that even though the most abundant lipids are similar in the different EV subtypes, there are differences in individual lipids and for some of the lipid classes, such as PE and Cer.

## 4. Discussion

Few studies have compared the proteome and lipidome of multiple EV subtypes. Most studies focus on a single EV subtype, limiting the knowledge of which proteins and lipids are common, and which are enriched in certain EV subpopulations. We present here an in-depth analysis of the proteome and lipidome of four subpopulations of EVs from immune cells to address this knowledge gap. The proteomes of the four EV subtypes were well separated and had distinct protein profiles. Mitochondrial and cytoskeleton proteins were enriched in L-EVs LD and L-EVs HD, respectively. Tetraspanins, ESCRT machinery proteins, and 14-3-3 proteins were enriched in S-EVs LD, while histones, complement factors, and CCT proteins were enriched in S-EVs HD. Proteins such as flotillins, RABs, heat shock proteins, guanine nucleotide-binding proteins (GNAs), and integrins were enriched in two or more EV subtypes. The lipidome was relatively similar for the EV subpopulations with lipids such as SM(34:1;O2) and PC(16:0_18:1), being the most abundant lipids in several of the EV subtypes. However, PE was substantially lower in L-EVs LD as compared to all the other subpopulations and ceramides were more abundant in L-EVs as compared to S-EVs.

The proteins enriched in L-EVs LD were associated with mitochondrion and endoplasmic reticulum. Specifically, proteins such as TIMM/TOMM, MICOS, and ATP-synthase proteins were significantly enriched in L-EVs fractions, but commonly with a higher enrichment in the L-EVs LD. This validates previous studies by us and others that L-EVs from both cell lines and clinical samples are enriched in mitochondrial proteins such as mitofilin (IMMT), ATP5O, ATP5F1B, and ATP5F1A (11, 28–35). Additionally, there are papers suggesting that L-EVs can transport whole mitochondria (36, 37). However, there are also studies showing that S-EVs contain, for example, mitochondrial DNA or mitochondrial proteins (12, 36, 38, 39). Furthermore, it has been suggested that mitochondria protein-containing vesicles, mitovesicles, are a novel population of EVs, distinct from both S-EVs and L-EVs such as exosomes and microvesicles (40). Lastly, it has also been demonstrated that the amount of EVs containing mitochondrial proteins is altered in Down syndrome, asthma and cancer (39–41). Puhm and colleagues also showed that stimulating THP-1 cells with LPS, the same cells that we used here, increased TOMM22 in L-EVs, suggesting that stress can enrich the mitochondrial content in L-EVs (42). Together, these findings suggest that there is an interaction or overlap between processes in the mitochondrion and the biogenesis of certain subpopulations of EVs and that this may be altered in pathophysiological conditions.

On the other hand, proteins enriched in L-EVs HD were associated with the cytosol, the cytoskeleton and the kinesin complex. Specifically, proteins associated with cytokinesis, such as microtubules- and central spindle complex-associated proteins, including KIFs, PRC1 and RACGAP1 were enriched. During cell division, RACGAP1 interacts with KIF23 and PRC1 to form and stabilize the central spindle, and it has been shown that RACGAP1 is overexpressed in several cancers (43). About half of the proteins belonging to the kinesin superfamily proteins (KIFs) were quantified in our proteomics analysis, and the majority of the quantified KIFs showed significant enrichment in L-EV HD. The enrichment observed in L-EVs HD isolated in this study validates previous studies published by us and others, where it was shown that KIFs and RACGAP1 are enriched in L-EVs (11, 31, 35, 44, 45) and further enriched in L-EVs HD (46). Rai *et al* suggested that the EVs containing the midbody remnants is a distinct EV subpopulation from exosomes (S-EVs) and microvesicles (L-EVs) and our data may support this as these proteins were uniquely upregulated in L-EV HD in both cell lines.

Additionally, some protein groups were enriched in both L-EVs LD and HD, such as heat shock proteins and voltage-dependent anion channels (VDAC). Heat shock proteins are involved in the folding, unfolding and stabilization of proteins and thereby ensure correct folding, but they also have a multitude of other functions. In our previous study, we found that the majority of heat shock proteins were enriched in L-EVs (11), which also others have observed (29, 33, 35, 47, 48). We could confirm it here, and further show that there was little difference between the L-EVs LD and HD. However, this time we also included the type two chaperonins “chaperonin containing TCP-1” (CCT) proteins which fold about 10% of all proteins with a focus on actin and tubulin, but also proteins essential for cancer development (49). Interestingly, these CCTs tended to be enriched in S-EVs instead, with the highest abundance in S-EVs HD, showing a distinct distribution compared to the other heat shock proteins. In previous studies, CCTs have been identified as both enriched in S-EVs (48) and L-EVs (35, 50). Interestingly, Rojas-Gómez and colleagues recently suggested that CCTs control EV production by affecting the number and size of lipid droplets, and by affecting the kinesin dynamics, and inter-organelle contacts and movement. Specifically, cells treated with siRNA to reduce the CCTs expression released more S-EVs, which was suggested to be an effect of the redirection of the endolysosomal pathway to multivesicular body biogenesis and EV release (51).

Additionally, some protein groups were enriched in both L-EVs HD and S-EVs HD, such as tubulins. Tubulins have previously been shown to be enriched in both L-EVs (35, 48) and S-EVs HD (52) demonstrating their presence in both L-EVs and S-EVs. Our data clearly showed that when separated based on density the tubulins would end up in the HD fraction for both L-EVs and S-EVs.

Besides the tubulins and CCTs mentioned above, S-EVs HD were also enriched in histones, complement proteins, apolipoproteins and collagens compared to the other EV subpopulations. At first glance, this may seem to be a contamination. However, these EVs have been bottom-loaded on a density cushion, specifically designed with a “buffer” fraction (32%), positioned between the loading fraction (37.5%) and the collection fraction of the HD EVs (above 32%). This means that histones, complement proteins, apolipoproteins and collagens must have been attached to a lipid structure to move up to this density, as free protein would not float. In recent years, the investigations of a protein corona surrounding EVs have increased. The EV protein corona is a group of proteins that are attached to the vesicle surface after the vesicle is released. The reason for their binding can be due to, but not limited to, electrostatic interactions, proximity, protein aggregation and receptor/ligand binding. Proteins that are adsorbed to EVs creating the corona are, for example albumin, apolipoprotein, complement factors, cytokines and fibrinogen (53, 54). We currently do not know if these proteins are attached as a corona or loaded inside the S-EV HD. If we speculate that these proteins are derived from a corona, we currently did not determine whether it is a biological feature, hence, S-EVs attract more proteins than L-EVs or if it is a technical issue, such as, that more proteins being forced to attach to S-EVs due to the higher g forces need to pellet crude S-EVs as compared to L-EVs. However, it has been suggested that glycoproteins on EVs can bind some of the corona proteins (53) and interestingly, glypican-1, glypican-2, syndican-2 and syndecan-4 are enriched in S-EVs in our dataset. Additionally, we also do not know if the HD and LD S-EVs are two distinct S-EV subpopulations and therefore attract these proteins differently or if the EVs are the same but due to more protein corona one subtype ends up at a higher density. Future studies about protein corona versus co-isolation of contaminants with different EV subtypes are needed. Intrestingly, a recent study showed that histones were associated with the membrane of tetraspanins-positive S-EVs (55). Furthermore, they observed that histones were also present in CD63-positive intraluminal vesicles in multivesicular bodies, showing that the histones can be released associated with S-EVs and hence, not attached as a corona after the secretion.

Proteins enriched in S-EVs LD were tetraspanins, ESCRT complex proteins, 14-3-3 and ADAM10. As S-EVs are the most studied EV subtype, they are commonly known to be present in S-EVs. We and others have previously also shown that they are enriched in S-EVs as compared to L-EVs and/or further enriched in S-EVs LD as compared to S-EVs HD (11, 29–31, 33, 35, 44, 47, 48, 52, 56–59) from both cell lines and clinical samples. ADAM10 belongs to a family of transmembrane proteins that are involved in ectodomain shedding and cell adhesion. Interestingly, it has been shown that ADAM10 is associated with tetraspanins on cells and that some tetraspanins have been shown to regulate the ADAM10 surface expression levels and its activation for its cleavage of several substrates (25–27). Furthermore, it has been shown that extracellular vesicles contain active ADAM10 that can cleave its substrate (60). We show here that ADAM10 is mainly expressed on EVs that are also expressing tetraspanins. Together, this suggests a close relationship between ADAMs and tetraspanins that may be of interest for future EV studies to determine in more detail.

Lastly, we also identified some protein groups that were enriched in two or more EV subpopulations. While flotillins were enriched in all subpopulations, GNA proteins, integrins and annexins were enriched in both S-EV LD and L-EV LD and Syntenin-1 and syndecans were enriched in both S-EVs LD and S-EVs HD. As some studies have previously suggested that annexins are enriched in L-EVs while others have suggested that they are enriched in S-EVs (30, 47, 48, 57, 61), these proteins are most likely not good markers to distinguish EV subpopulations as their expression may be different depending on cell source. Most other studies have suggested that integrins are enriched in S-EVs (33, 44, 56, 61, 62) and we observe that for some of the integrins, while others are enriched in both S-EVs LD and L-EVs LD, which suggests that also their expressions are cell source dependent. Syntenin-1 (SDCBP) has mainly been shown previously to be enriched in S-EVs by us and others (11, 29, 35, 52, 56, 57, 63, 64) and has also been suggested to be part of the biogenesis of exosomes together with syndecans and ALIX (PDCD6IP) (65). We report here that we observed similar enrichment in S-EVs HD and LD for syntenin-1 and syndecans, which may suggest that both these EV types have similar biogenesis.

Next, we also determined the lipid content of the different subpopulations of EVs. In contrast to the proteomic analysis, which is unbiased, the lipidomic analysis is targeted, with a list of 107 lipids being included in our analysis. Cholesterol has been suggested to make up over 40% of EV lipids (66, 67). However, as cholesterol was not included in our analysis, this should be considered when comparing percentages between studies.

Overall, the lipid content of S-EVs and L-EVs was quite similar, which confirms previous observations by others in EVs from seminal plasma and cell lines (32, 64). Haraszti and colleagues, for example, observed that the lipid content of L-EVs and S-EVs was more similar than the protein content. However, when comparing EVs from three different cell lines they noted differences between the cell lines (32). Our findings were consistent with this observation. Durcin *et al* also suggested similarity in the lipid profile of L-EVs and S-EVs except for cholesterol content and externalised PS, two parameters we did not evaluate in our study (48). Zhang *et al* demonstrated that S-EVs of different sizes had similar lipid cargo, however, exomers had distinct lipids (68). Also, Zhang and coworkers showed that the lipid content in EVs was cell-type specific. However, we and others have detected differences in sphingolipids and phospholipids. For example, others have observed more Cer in L-EVs compared to S-EVs, while more SM was observed in S-EVs compared to L-EVs (69–72). This could be verified in our study as we identified more Cer in L-EVs in both cell lines and more SM in S-EVs in THP-1. The most profound difference in our dataset was that PE was lower in L-EVs LD compared to all three of the other subpopulations. Tacconi *et al* focused their analysis on how the lipid cargo of S-EVs and L-EVs from macrophages was altered in high glucose environments, but their data suggest that L-EVs have less PE (70). Additionally, S-EVs from platelets have been shown to have more PE than L-EVs (71).

In conclusion, our study shows that the proteome and lipidome of large and small EV subpopulations of different densities are substantially different. However, this was more prominent for the proteome than for the lipidome. Importantly, we cannot exclude that additional EV subtypes exist within these four subpopulations, and future studies will have to dissect this further. Here, we suggest proteins and lipids that are enriched in one or more of our EV subpopulations, but have also identified molecules that are similarly expressed in all EV subtypes.

**Supplementary Figure 1. LD and HD samples.** Representative photographs of density cushions for L-EVs and S-EVs from HMC-1.

Supplementary Figure 2. Particle and protein measurement of all samples. (A) Particles were measured with NTA. (B) Proteins were measured with

**Supplementary Figure 3. Higher magnification of TEM electrogram.** Representative micrographs are shown for the four EV subtypes from HMC-1 and THP-1. The scale bars represent 200 nm.

**Supplementary Figure 4. Pairwise comparisons of the proteome of the four subtypes of EVs.** (**A-D**) Volcano plots of the proteomes of S-EVs LD, S-EVs HD, L-EVs LD, and L-EVs HD from HMC-1 compared with each other. (**E**) Summary of the numbers of altered proteins in each comparison in A-D. (**F-I**) Volcano plots of the proteomes of S-EVs LD, S-EVs HD, L-EVs LD, and L-EVs HD from THP-1 compared with each other. (**J**) Summary of the numbers of altered proteins in each comparison in F-I. Dotted lines indicate cutoffs; 1.3 on the y-axis (corresponding to p < 0.05) and 0.585 on the x-axis (corresponding to fold change >1.5).

**Supplementary Figure 5. A step-by-step explanation of how the uniquely enriched proteins in each EV subpopulation for Figure 2D-G were calculated.**

**Supplementary Figure 6. The expression of MISEV guideline proteins in the four EV subpopulations from HMC-1 cells. (A-E)** In the MISEV guidelines (13, 24), five categories of proteins are recommended to be evaluated in EV isolates. We have analysed all proteins in our dataset that belong to these five categories. (**A**) Category 1 – Membrane proteins associated with the plasma membrane (PM)/endosomes. (**B**) Category 2 – Cytosolic proteins in EVs. (**C**) Category 3 – Major components of non-EV co-isolated structures. (**D**) Category 4 – Proteins associated with intracellular compartments other than plasma membrane (PM)/endosomes. (**E**) Category 5 – Secreted proteins recovered with EVs.

**Supplementary Figure 7. Triton-X controls for the nanoFCM experiments.**

**Supplementary Figure 8. Buffer and antibody controls for the nanoFCM experiments.**

**(A)** The buffers used in this project were analysed in the Flow NanoAnlayzer to determine the background. **(B)** Controls evaluating the background generated by unbound antibody, the autofluorescence by the EVs themselves, and the isotype control for the antibodies were evaluated.

**Supplementary Figure 9. Controls for double staining for the nanoFCM experiments.** To determine that the double staining of the EVs for both ADAM10 and the tetraspanins did not affect the percentage of positive EVs for any of the markers, we compared single and double staining.

**Supplementary Table 1. List of the 107 lipids that we analysed in this study and how they were identified.**

## Acknowledgement

We thank the Centre for Cellular Imaging at the University of Gothenburg and the National Microscopy Infrastructure (VR-RFI 2016-00968) for microscopy support. We also wish to acknowledge Annika Thorsell and Elham Rekabdar at the Proteomics Core Facility at Sahlgrenska Academy, University of Gothenburg, for performing the LC-tandem MS analysis. The Proteomics Core Facility, Sahlgrenska academy, Gothenburg University, is supported with financial support from SciLifeLab and BioMS.

## 6. Authors’ contributions202

**Anna Lischnig**; methodology, formal analysis, investigation, writing original draft, visualization. **Nasibeh Karimi**; methodology, investigation, writing – review and editing. **Per Larsson**; methodology, formal analysis, investigation, writing – review and editing. **Karin Ekström**; methodology, formal analysis, investigation, writing -review and editing. **Rossella Crescitelli**; methodology, investigation, writing - review and editing. **Anna-Carin Olin**; investigation, writing - review and editing. **Cecilia Lässer**; conceptualization, methodology, formal analysis, investigation, writing - original draft, visualization, supervision, project administration, funding acquisition.

## References

1. Couch Y, Buzas EI, Di Vizio D, Gho YS, Harrison P, Hill AF, et al. A brief history of nearly EV-erything - The rise and rise of extracellular vesicles. J Extracell Vesicles. 2021;10(14):e12144.

2. Yates AG, Pink RC, Erdbrugger U, Siljander PR, Dellar ER, Pantazi P, et al. In sickness and in health: The functional role of extracellular vesicles in physiology and pathology in vivo: Part I: Health and Normal Physiology: Part I: Health and Normal Physiology. J Extracell Vesicles. 2022;11(1):e12151.

3. Yates AG, Pink RC, Erdbrugger U, Siljander PR, Dellar ER, Pantazi P, et al. In sickness and in health: The functional role of extracellular vesicles in physiology and pathology in vivo: Part II: Pathology: Part II: Pathology. J Extracell Vesicles. 2022;11(1):e12190.

4. Lasser C, Jang SC, Lotvall J. Subpopulations of extracellular vesicles and their therapeutic potential. Mol Aspects Med. 2018;60:1–14.

5. Wiklander OPB, Brennan MA, Lotvall J, Breakefield XO, El Andaloussi S. Advances in therapeutic applications of extracellular vesicles. Sci Transl Med. 2019;11(492).

6. Phillips W, Willms E, Hill AF. Understanding extracellular vesicle and nanoparticle heterogeneity: Novel methods and considerations. Proteomics. 2021;21(13-14):e2000118.

7. Fernandis AZ, Wenk MR. Membrane lipids as signaling molecules. Curr Opin Lipidol. 2007;18(2):121–8.

8. Butterfield JH, Weiler D, Dewald G, Gleich GJ. Establishment of an immature mast cell line from a patient with mast cell leukemia. Leuk Res. 1988;12(4):345–55.

9. Nilsson G, Blom T, Kusche-Gullberg M, Kjellen L, Butterfield JH, Sundstrom C, et al. Phenotypic characterization of the human mast-cell line HMC-1. Scandinavian journal of immunology. 1994;39(5):489–98.

10. Tsuchiya S, Yamabe M, Yamaguchi Y, Kobayashi Y, Konno T, Tada K. Establishment and characterization of a human acute monocytic leukemia cell line (THP-1). Int J Cancer. 1980;26(2):171–6.

11. Lischnig A, Bergqvist M, Ochiya T, Lasser C. Quantitative Proteomics Identifies Proteins Enriched in Large and Small Extracellular Vesicles. Mol Cell Proteomics. 2022;21(9):100273.

12. Lazaro-Ibanez E, Lasser C, Shelke GV, Crescitelli R, Jang SC, Cvjetkovic A, et al. DNA analysis of low- and high-density fractions defines heterogeneous subpopulations of small extracellular vesicles based on their DNA cargo and topology. J Extracell Vesicles. 2019;8(1):1656993.

13. Welsh JA, Goberdhan DCI, O’Driscoll L, Buzas EI, Blenkiron C, Bussolati B, et al. Minimal information for studies of extracellular vesicles (MISEV2023): From basic to advanced approaches. J Extracell Vesicles. 2024;13(2):e12404.

14. Lim HJ, Yoon H, Kim H, Kang YW, Kim JE, Kim OY, et al. Extracellular Vesicle Proteomes Shed Light on the Evolutionary, Interactive, and Functional Divergence of Their Biogenesis Mechanisms. Front Cell Dev Biol. 2021;9:734950.

15. Bergqvist M, Park KS, Karimi N, Yu L, Lasser C, Lotvall J. Extracellular vesicle surface engineering with integrins (ITGAL & ITGB2) to specifically target ICAM-1-expressing endothelial cells. J Nanobiotechnology. 2025;23(1):64.

16. Bergqvist M, Lasser C, Crescitelli R, Park KS, Lotvall J. A Non-Centrifugation Method to Concentrate and Purify Extracellular Vesicles Using Superabsorbent Polymer Followed by Size Exclusion Chromatography. J Extracell Vesicles. 2025;14(1):e70037.

17. Welsh JA, Van Der Pol E, Arkesteijn GJA, Bremer M, Brisson A, Coumans F, et al. MIFlowCyt-EV: a framework for standardized reporting of extracellular vesicle flow cytometry experiments. J Extracell Vesicles. 2020;9(1):1713526.

18. Welsh JA, Arkesteijn GJA, Bremer M, Cimorelli M, Dignat-George F, Giebel B, et al. A compendium of single extracellular vesicle flow cytometry. J Extracell Vesicles. 2023;12(2):e12299.

19. Osteikoetxea X, Sodar B, Nemeth A, Szabo-Taylor K, Paloczi K, Vukman KV, et al. Differential detergent sensitivity of extracellular vesicle subpopulations. Org Biomol Chem. 2015;13(38):9775–82.

20. Deutsch EW, Bandeira N, Sharma V, Perez-Riverol Y, Carver JJ, Kundu DJ, et al. The ProteomeXchange consortium in 2020: enabling ’big data’ approaches in proteomics. Nucleic acids research. 2020;48(D1):D1145–D52.

21. Huang da W, Sherman BT, Lempicki RA. Bioinformatics enrichment tools: paths toward the comprehensive functional analysis of large gene lists. Nucleic acids research. 2009;37(1):1–13.

22. Consortium E-T, Van Deun J, Mestdagh P, Agostinis P, Akay O, Anand S, et al. EV-TRACK: transparent reporting and centralizing knowledge in extracellular vesicle research. Nature methods. 2017;14(3):228–32.

23. Lotvall J, Hill AF, Hochberg F, Buzas EI, Di Vizio D, Gardiner C, et al. Minimal experimental requirements for definition of extracellular vesicles and their functions: a position statement from the International Society for Extracellular Vesicles. J Extracell Vesicles. 2014;3:26913.

24. Thery C, Witwer KW, Aikawa E, Alcaraz MJ, Anderson JD, Andriantsitohaina R, et al. Minimal information for studies of extracellular vesicles 2018 (MISEV2018): a position statement of the International Society for Extracellular Vesicles and update of the MISEV2014 guidelines. J Extracell Vesicles. 2018;7(1):1535750.

25. Arduise C, Abache T, Li L, Billard M, Chabanon A, Ludwig A, et al. Tetraspanins regulate ADAM10-mediated cleavage of TNF-alpha and epidermal growth factor. J Immunol. 2008;181(10):7002–13.

26. Yanez-Mo M, Sanchez-Madrid F, Cabanas C. Membrane proteases and tetraspanins. Biochem Soc Trans. 2011;39(2):541–6.

27. Jouannet S, Saint-Pol J, Fernandez L, Nguyen V, Charrin S, Boucheix C, et al. TspanC8 tetraspanins differentially regulate the cleavage of ADAM10 substrates, Notch activation and ADAM10 membrane compartmentalization. Cellular and molecular life sciences : CMLS. 2016;73(9):1895–915.

28. Wu L, Zhou L, An J, Shao X, Zhang H, Wang C, et al. Comprehensive profiling of extracellular vesicles in uveitis and scleritis enables biomarker discovery and mechanism exploration. J Transl Med. 2023;21(1):388.

29. Rai A, Fang H, Claridge B, Simpson RJ, Greening DW. Proteomic dissection of large extracellular vesicle surfaceome unravels interactive surface platform. J Extracell Vesicles. 2021;10(13):e12164.

30. Kowal J, Arras G, Colombo M, Jouve M, Morath JP, Primdal-Bengtson B, et al. Proteomic comparison defines novel markers to characterize heterogeneous populations of extracellular vesicle subtypes. Proc Natl Acad Sci U S A. 2016;113(8):E968–77.

31. Dozio V, Sanchez JC. Characterisation of extracellular vesicle-subsets derived from brain endothelial cells and analysis of their protein cargo modulation after TNF exposure. J Extracell Vesicles. 2017;6(1):1302705.

32. Haraszti RA, Didiot MC, Sapp E, Leszyk J, Shaffer SA, Rockwell HE, et al. High-resolution proteomic and lipidomic analysis of exosomes and microvesicles from different cell sources. J Extracell Vesicles. 2016;5:32570.

33. Minciacchi VR, You S, Spinelli C, Morley S, Zandian M, Aspuria PJ, et al. Large oncosomes contain distinct protein cargo and represent a separate functional class of tumor-derived extracellular vesicles. Oncotarget. 2015;6(13):11327–41.

34. Ahmadzada T, Vijayan A, Vafaee F, Azimi A, Reid G, Clarke S, et al. Small and Large Extracellular Vesicles Derived from Pleural Mesothelioma Cell Lines Offer Biomarker Potential. Cancers (Basel). 2023;15(8).

35. Xu R, Greening DW, Rai A, Ji H, Simpson RJ. Highly-purified exosomes and shed microvesicles isolated from the human colon cancer cell line LIM1863 by sequential centrifugal ultrafiltration are biochemically and functionally distinct. Methods. 2015;87:11–25.

36. Phinney DG, Di Giuseppe M, Njah J, Sala E, Shiva S, St Croix CM, et al. Mesenchymal stem cells use extracellular vesicles to outsource mitophagy and shuttle microRNAs. Nature communications. 2015;6:8472.

37. D’Souza A, Burch A, Dave KM, Sreeram A, Reynolds MJ, Dobbins DX, et al. Microvesicles transfer mitochondria and increase mitochondrial function in brain endothelial cells. J Control Release. 2021;338:505–26.

38. Rabas N, Palmer S, Mitchell L, Ismail S, Gohlke A, Riley JS, et al. PINK1 drives production of mtDNA-containing extracellular vesicles to promote invasiveness. The Journal of cell biology. 2021;220(12).

39. Hough KP, Trevor JL, Strenkowski JG, Wang Y, Chacko BK, Tousif S, et al. Exosomal transfer of mitochondria from airway myeloid-derived regulatory cells to T cells. Redox Biol. 2018;18:54–64.

40. D’Acunzo P, Perez-Gonzalez R, Kim Y, Hargash T, Miller C, Alldred MJ, et al. Mitovesicles are a novel population of extracellular vesicles of mitochondrial origin altered in Down syndrome. Sci Adv. 2021;7(7).

41. Jang SC, Crescitelli R, Cvjetkovic A, Belgrano V, Olofsson Bagge R, Sundfeldt K, et al. Mitochondrial protein enriched extracellular vesicles discovered in human melanoma tissues can be detected in patient plasma. J Extracell Vesicles. 2019;8(1):1635420.

42. Puhm F, Afonyushkin T, Resch U, Obermayer G, Rohde M, Penz T, et al. Mitochondria Are a Subset of Extracellular Vesicles Released by Activated Monocytes and Induce Type I IFN and TNF Responses in Endothelial Cells. Circulation research. 2019;125(1):43–52.

43. Eid RA, Soltan MA, Eldeen MA, Shati AA, Dawood SA, Eissa M, et al. Assessment of RACGAP1 as a Prognostic and Immunological Biomarker in Multiple Human Tumors: A Multiomics Analysis. Int J Mol Sci. 2022;23(22).

44. Martin-Jaular L, Nevo N, Schessner JP, Tkach M, Jouve M, Dingli F, et al. Unbiased proteomic profiling of host cell extracellular vesicle composition and dynamics upon HIV-1 infection. EMBO J. 2021;40(8):e105492.

45. Cocozza F, Martin-Jaular L, Lippens L, Di Cicco A, Arribas YA, Ansart N, et al. Extracellular vesicles and co-isolated endogenous retroviruses from murine cancer cells differentially affect dendritic cells. EMBO J. 2023;42(24):e113590.

46. Rai A, Greening DW, Xu R, Chen M, Suwakulsiri W, Simpson RJ. Secreted midbody remnants are a class of extracellular vesicles molecularly distinct from exosomes and microparticles. Commun Biol. 2021;4(1):400.

47. Pfeiffer A, Petersen JD, Falduto GH, Anderson DE, Zimmerberg J, Metcalfe DD, et al. Selective immunocapture reveals neoplastic human mast cells secrete distinct microvesicle-and exosome-like populations of KIT-containing extracellular vesicles. J Extracell Vesicles. 2022;11(10):e12272.

48. Durcin M, Fleury A, Taillebois E, Hilairet G, Krupova Z, Henry C, et al. Characterisation of adipocyte-derived extracellular vesicle subtypes identifies distinct protein and lipid signatures for large and small extracellular vesicles. J Extracell Vesicles. 2017;6(1):1305677.

49. Showalter AE, Martini AC, Nierenberg D, Hosang K, Fahmi NA, Gopalan P, et al. Investigating Chaperonin-Containing TCP-1 subunit 2 as an essential component of the chaperonin complex for tumorigenesis. Scientific reports. 2020;10(1):798.

50. Mariscal J, Vagner T, Kim M, Zhou B, Chin A, Zandian M, et al. Comprehensive palmitoyl-proteomic analysis identifies distinct protein signatures for large and small cancer-derived extracellular vesicles. J Extracell Vesicles. 2020;9(1):1764192.

51. Rojas-Gomez A, Dosil SG, Chichon FJ, Fernandez-Gallego N, Ferrarini A, Calvo E, et al. Chaperonin CCT controls extracellular vesicle production and cell metabolism through kinesin dynamics. J Extracell Vesicles. 2023;12(6):e12333.

52. Jeppesen DK, Fenix AM, Franklin JL, Higginbotham JN, Zhang Q, Zimmerman LJ, et al. Reassessment of Exosome Composition. Cell. 2019;177(2):428–45 e18.

53. Buzas EI. Opportunities and challenges in studying the extracellular vesicle corona. Nat Cell Biol. 2022;24(9):1322–5.

54. Heidarzadeh M, Zarebkohan A, Rahbarghazi R, Sokullu E. Protein corona and exosomes: new challenges and prospects. Cell communication and signaling : CCS. 2023;21(1):64.

55. Singh B, Fredriksson Sundbom M, Muthukrishnan U, Natarajan B, Stransky S, Gorgens A, et al. Extracellular Histones as Exosome Membrane Proteins Regulated by Cell Stress. J Extracell Vesicles. 2025;14(2):e70042.

56. Yokoi A, Ukai M, Yasui T, Inokuma Y, Hyeon-Deuk K, Matsuzaki J, et al. Identifying high-grade serous ovarian carcinoma-specific extracellular vesicles by polyketone-coated nanowires. Sci Adv. 2023;9(27):eade6958.

57. Panizza E, Regalado BD, Wang F, Nakano I, Vacanti NM, Cerione RA, et al. Proteomic analysis reveals microvesicles containing NAMPT as mediators of radioresistance in glioma. Life Sci Alliance. 2023;6(6).

58. Li D, Lai W, Wang Q, Xiang Z, Nan X, Yang X, et al. CD151 enrichment in exosomes of luminal androgen receptor breast cancer cell line contributes to cell invasion. Biochimie. 2021;189:65–75.

59. Keerthikumar S, Gangoda L, Liem M, Fonseka P, Atukorala I, Ozcitti C, et al. Proteogenomic analysis reveals exosomes are more oncogenic than ectosomes. Oncotarget. 2015;6(17):15375–96.

60. Stoeck A, Keller S, Riedle S, Sanderson MP, Runz S, Le Naour F, et al. A role for exosomes in the constitutive and stimulus-induced ectodomain cleavage of L1 and CD44. The Biochemical journal. 2006;393(Pt 3):609–18.

61. Turner DL, Korneev DV, Purdy JG, de Marco A, Mathias RA. The host exosome pathway underpins biogenesis of the human cytomegalovirus virion. Elife. 2020;9.

62. Graca AL, Gomez-Florit M, Osorio H, Rodrigues MT, Domingues RMA, Reis RL, et al. Controlling the fate of regenerative cells with engineered platelet-derived extracellular vesicles. Nanoscale. 2022;14(17):6543–56.

63. Kugeratski FG, Hodge K, Lilla S, McAndrews KM, Zhou X, Hwang RF, et al. Quantitative proteomics identifies the core proteome of exosomes with syntenin-1 as the highest abundant protein and a putative universal biomarker. Nat Cell Biol. 2021;23(6):631–41.

64. Chiasserini D, Mazzoni M, Bordi F, Sennato S, Susta F, Orvietani PL, et al. Identification and Partial Characterization of Two Populations of Prostasomes by a Combination of Dynamic Light Scattering and Proteomic Analysis. J Membr Biol. 2015;248(6):991–1004.

65. Baietti MF, Zhang Z, Mortier E, Melchior A, Degeest G, Geeraerts A, et al. Syndecan-syntenin-ALIX regulates the biogenesis of exosomes. Nat Cell Biol. 2012;14(7):677–85.

66. Skotland T, Sagini K, Sandvig K, Llorente A. An emerging focus on lipids in extracellular vesicles. Adv Drug Deliv Rev. 2020;159:308–21.

67. Brouwers JF, Aalberts M, Jansen JW, van Niel G, Wauben MH, Stout TA, et al. Distinct lipid compositions of two types of human prostasomes. Proteomics. 2013(Epub ahead of print).

68. Zhang H, Freitas D, Kim HS, Fabijanic K, Li Z, Chen H, et al. Identification of distinct nanoparticles and subsets of extracellular vesicles by asymmetric flow field-flow fractionation. Nat Cell Biol. 2018;20(3):332–43.

69. Martinez-Diaz P, Parra A, Sanchez-Lopez CM, Casas J, Lucas X, Marcilla A, et al. Small and Large Extracellular Vesicles of Porcine Seminal Plasma Differ in Lipid Profile. Int J Mol Sci. 2024;25(13).

70. Tacconi S, Vari F, Sbarigia C, Vardanyan D, Longo S, Mura F, et al. M1-derived extracellular vesicles polarize recipient macrophages into M2-like macrophages and alter skeletal muscle homeostasis in a hyper-glucose environment. Cell communication and signaling : CCS. 2024;22(1):193.

71. De Paoli SH, Tegegn TZ, Elhelu OK, Strader MB, Patel M, Diduch LL, et al. Dissecting the biochemical architecture and morphological release pathways of the human platelet extracellular vesiculome. Cellular and molecular life sciences : CMLS. 2018;75(20):3781–801.

72. Luo P, Mao K, Xu J, Wu F, Wang X, Wang S, et al. Metabolic characteristics of large and small extracellular vesicles from pleural effusion reveal biomarker candidates for the diagnosis of tuberculosis and malignancy. J Extracell Vesicles. 2020;9(1):1790158.

73. Kim D-K, Kang B, Kim OY, Choi D-s, Lee J, Kim SR, et al. EVpedia: an integrated database of high-throughput data for systemic analyses of extracellular vesicles. Journal of Extracellular Vesicles. 2013;2.

74. Chitti SV, Gummadi S, Kang T, Shahi S, Marzan AL, Nedeva C, et al. Vesiclepedia 2024: an extracellular vesicles and extracellular particles repository. Nucleic acids research. 2024;52(D1):D1694–D8.

