## Supplementary Figures and Table for "Proteomic and lipidomic profiling of immune cell-derived subpopulations of extracellular vesicles"

### Supplementary Figure 1

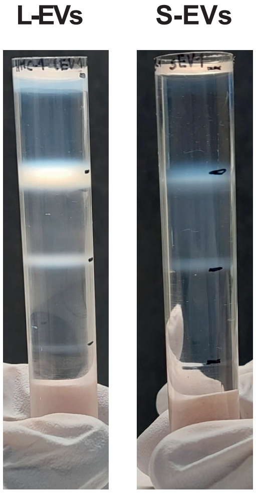

### Supplementary Figure 2

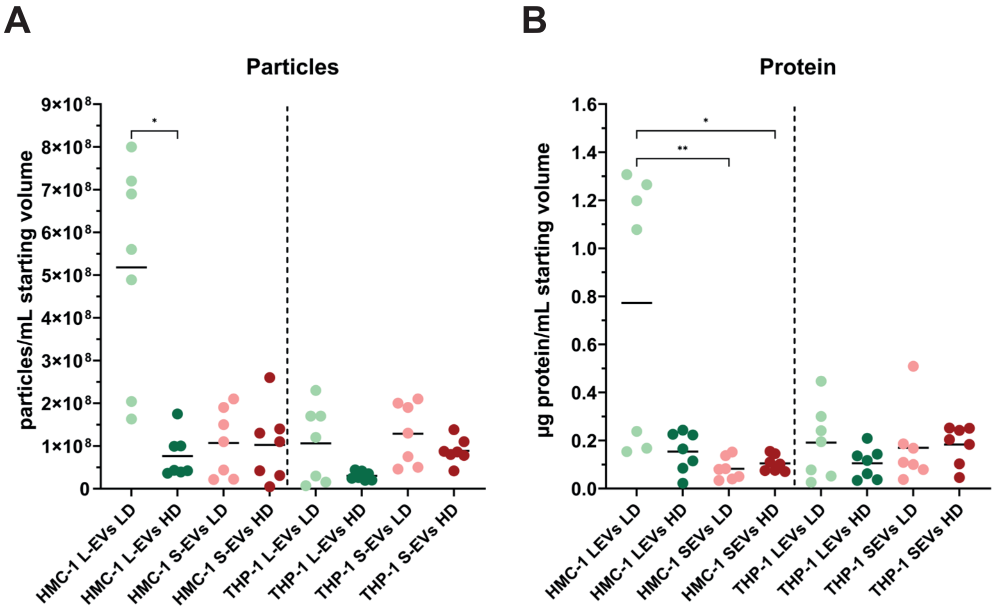

### Supplementary Figure 3

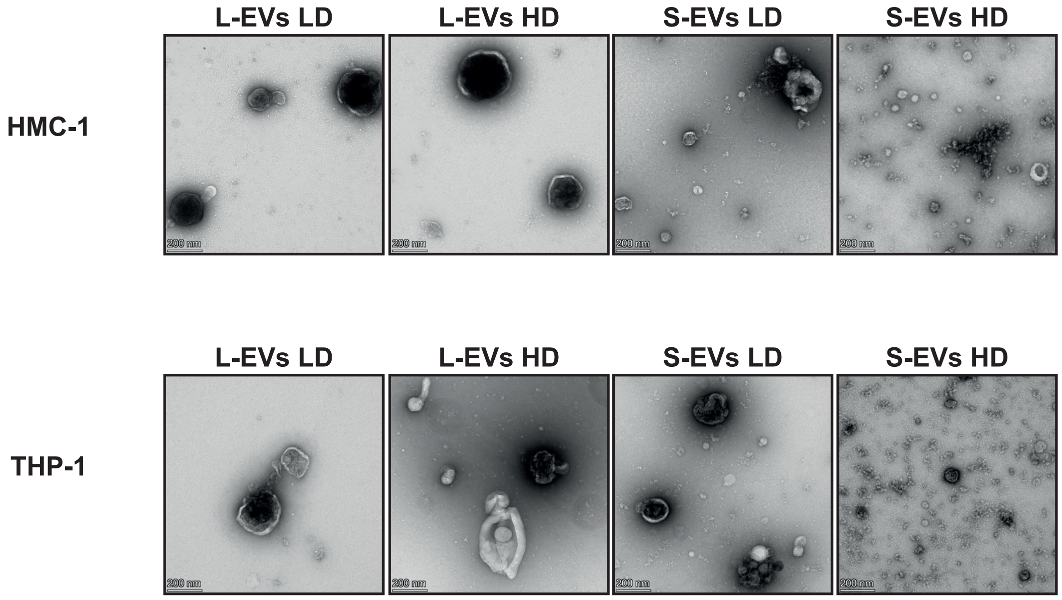

### Supplementary Figure 4

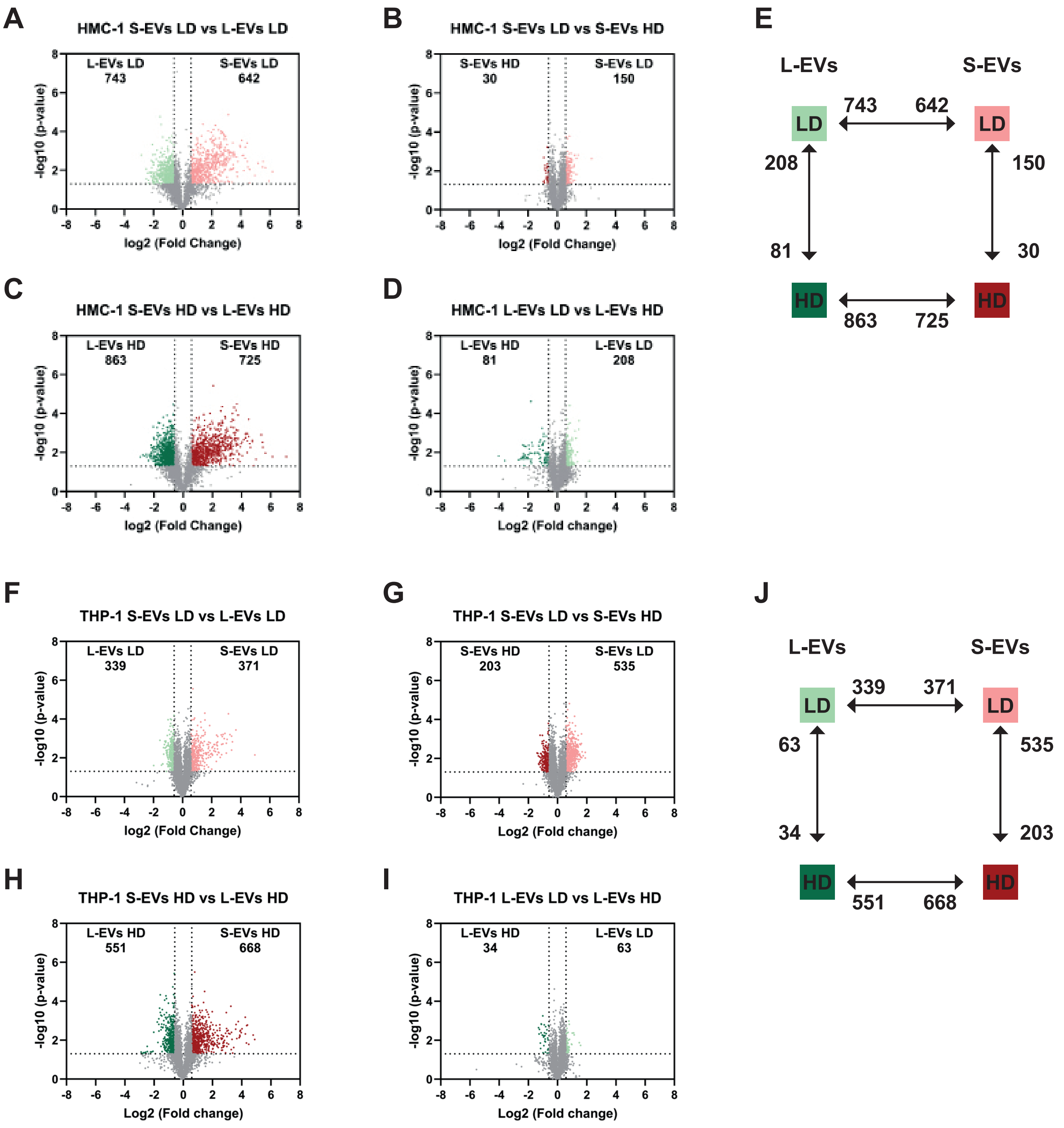

### Supplementary Figure 5

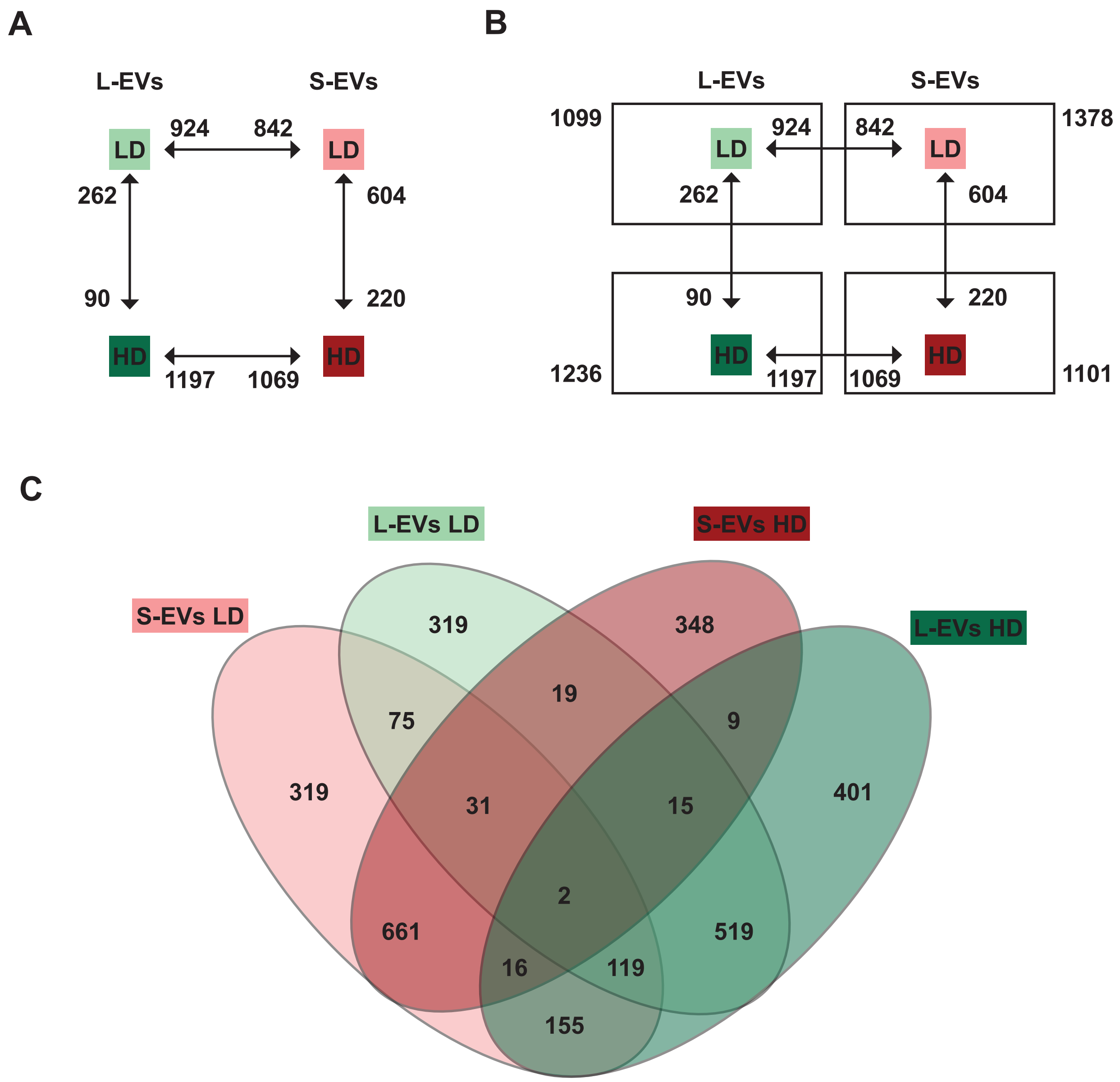


**D**

**Step 1.** Created volcano plots by pairwise comparisons between EV subtypes for each cell type. The comparisons that were made was:

L-EVs LD vs S-EVs LD (Supplementary Figure 4A and F),
S-EVs LD vs S-EVs HD (Supplementary Figure 4B and G),
S-EVs HD vs L-EVs HD (Supplementary Figure 4C and H),
L-EVs HD vs L-EVs LD (Supplementary Figure 4D and I)

They are summarised per cell type in Supplementary Figure 4E and J.

**Step 2.** Combined the lists of altered proteins from step 1 for both cell lines (Supplementary Figure 5A).

**Step 3.** Combined the lists of altered proteins from step 2 per EV subtype (Supplementary Figure 5B).

**Step 4.** Compared the lists form Step 3 to identify the proteins only altered in one subtype of EVs (Supplementary Figure 5C).

**Step 5.** Use the uniquely altered proteins in each EV subtype to be analysed in DAVID (Figure 2D-G).

### Supplementary Figure 6

**A**

**Category 1**
Membrane proteins associated with PM/ endosomes

**B**

**Category 2**
Cytosolic proteins in EVs

**C**

**Category 3**
Major components of non-EV co-isolated structures

**D**

**Category 4**
Proteins associated with intracellular compartments other than PM/endosomes

**E**

**Category 5**
Secreted proteins recovered with EVs

**HMC-1**

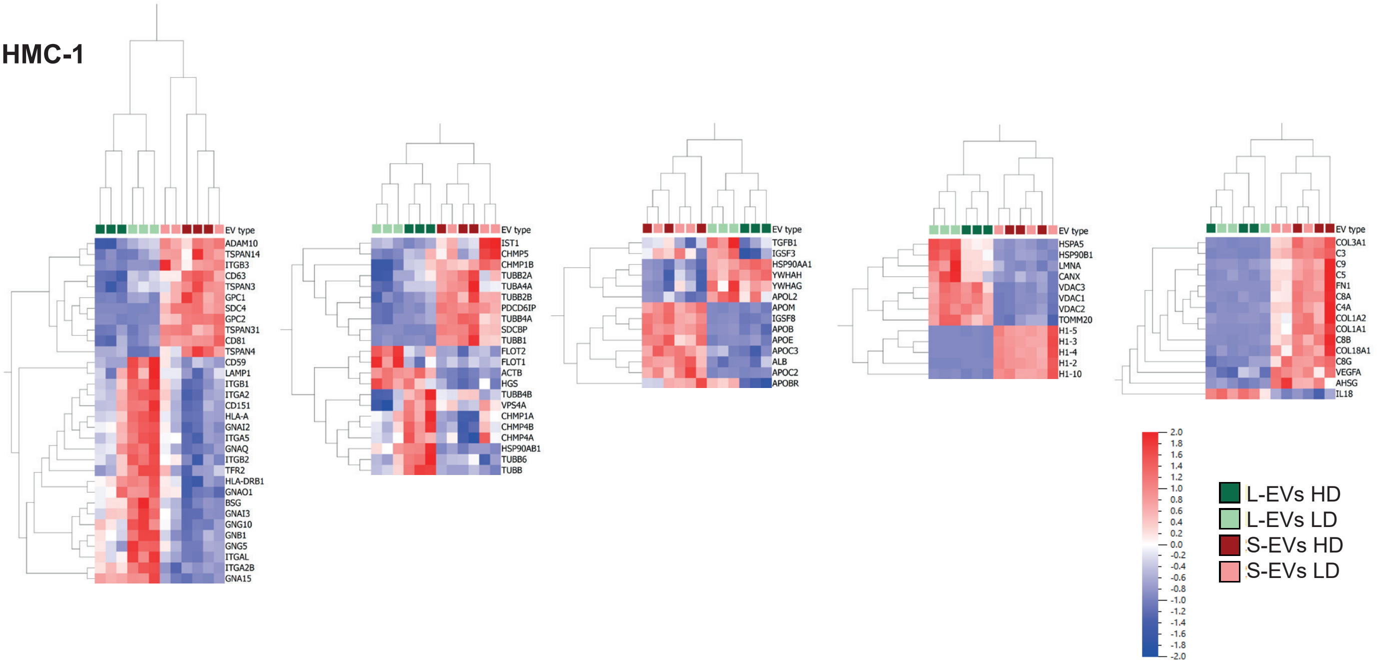

### Supplementary Figure 7

#### S-EVs

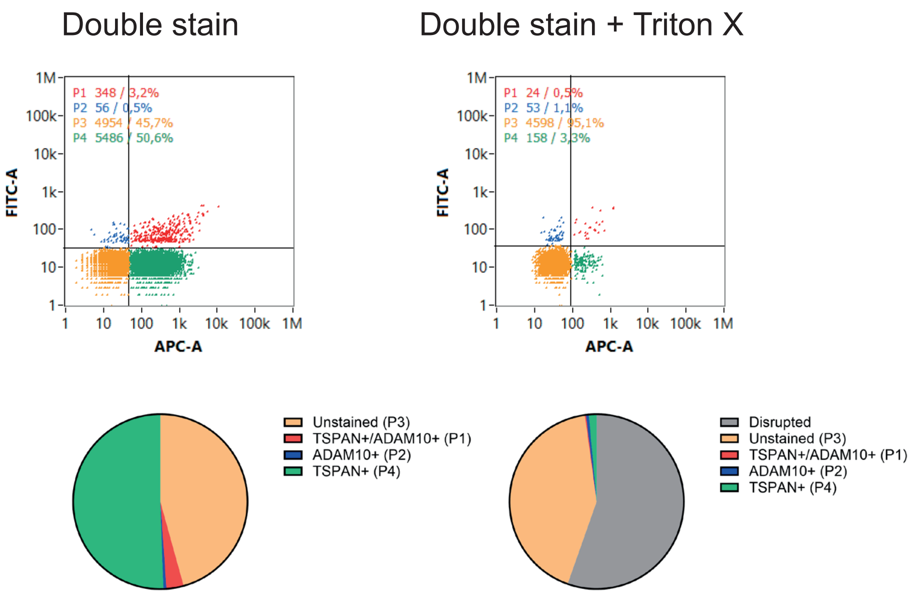

### Supplementary Figure 8

#### Buffer only controls

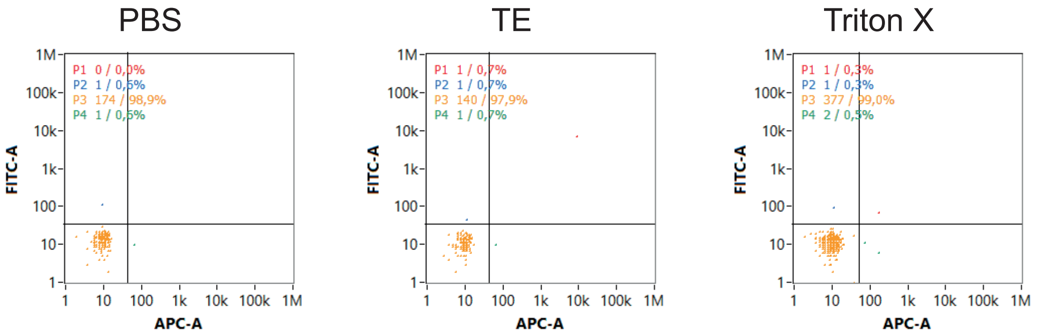


#### Antibody controls

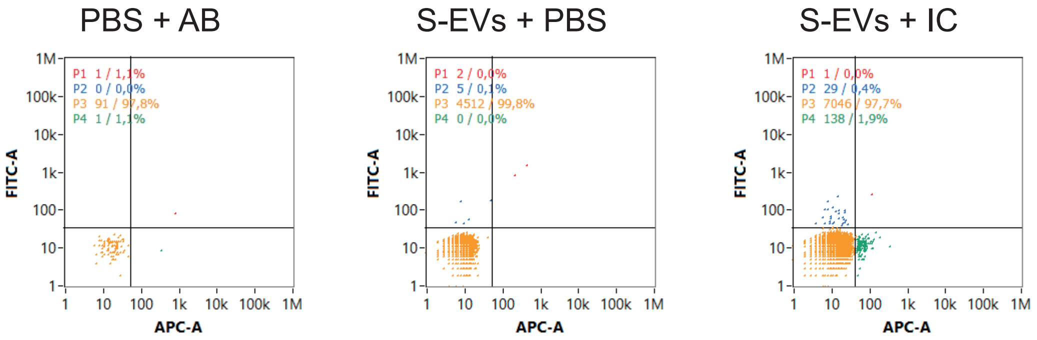

### Supplementary Figure 9

#### S-EVs

Single stain FITC

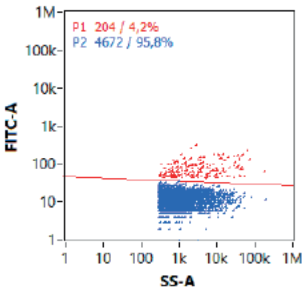


Double stain FITC

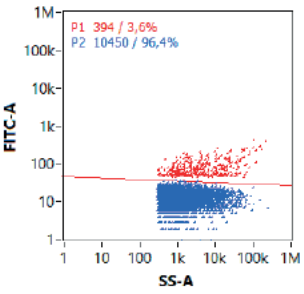


Single stain APC

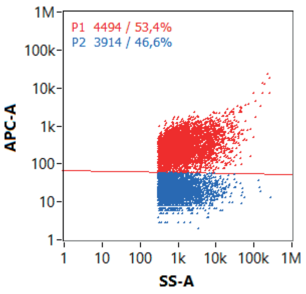


Double stain APC

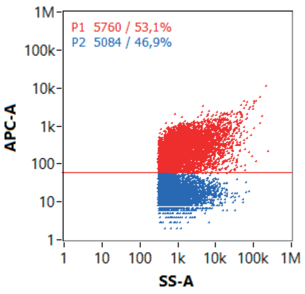

| Lipid_Class | Lipid_Annotation | Peak_Annotation_In_Method | Chromatography Columns_SKU HILIC:Waters_186009508 RP:Waters_186010360 | Retention Time Avg (min) | Precursor Ion | MS1 *(m/z)* | MS2 *(m/z)* | Internal Standard (Avanti_SPLASH 330707) |
|---|---|---|---|---|---|---|---|---|
| HexCer_SIL | GlcCer(d18:1(d5)/18:0) | xHexCer-d18_1-18_0-269-1 | RP_2* | 2,4 | [M+H]1+ | 733,6 | 269,3 | Stable Isotope Standard |
| Cer | Cer(d18:1/14:0) | Cer-d18_1-14_0-264-1 | RP_2 | 1,9 | [M+H]1+ | 510,5 | 264,3 | GlcCer(d18:1(d5)/18:0) |
| Cer | Cer(d18:1/16:0) | Cer-d18_1-16_0-264-1 | RP_2 | 2,4 | [M+H]1+ | 538,5 | 264,3 | GlcCer(d18:1(d5)/18:0) |
| Cer | Cer(d18:1/16:1) | Cer-d18_1-16_1-264-1 | RP_2 | 2,1 | [M+H]1+ | 536,5 | 264,3 | GlcCer(d18:1(d5)/18:0) |
| Cer | Cer(d18:1/17:0) | Cer-d18_1-17_0-264-1 | RP_2 | 2,7 | [M+H]1+ | 552,5 | 264,3 | GlcCer(d18:1(d5)/18:0) |
| Cer | Cer(d18:1/18:0) | Cer-d18_1-18_0-264-1 | RP_2 | 3,0 | [M+H]1+ | 566,6 | 264,3 | GlcCer(d18:1(d5)/18:0) |
| Cer | Cer(d18:1/20:0) | Cer-d18_1-20_0-264-1 | RP_2 | 3,7 | [M+H]1+ | 594,6 | 264,3 | GlcCer(d18:1(d5)/18:0) |
| Cer | Cer(d18:1/20:4) | Cer-d18_1-20_4-264-1 | RP_2 | 4,6 | [M+H]1+ | 586,5 | 264,3 | GlcCer(d18:1(d5)/18:0) |
| Cer | Cer(d18:1/22:0) | Cer-d18_1-22_0-264-1 | RP_2 | 4,6 | [M+H]1+ | 622,6 | 264,3 | GlcCer(d18:1(d5)/18:0) |
| Cer | Cer(d18:1/22:2) | Cer-d18_1-22_2-264-1 | RP_2 | 4,9 | [M+H]1+ | 618,6 | 264,3 | GlcCer(d18:1(d5)/18:0) |
| Cer | Cer(d18:1/22:3) | Cer-d18_1-22_3-264-1 | RP_2 | 4,2 | [M+H]1+ | 616,6 | 264,3 | GlcCer(d18:1(d5)/18:0) |
| Cer | Cer(d18:1/22:4) | Cer-d18_1-22_4-264-1 | RP_2 | 5,3 | [M+H]1+ | 614,6 | 264,3 | GlcCer(d18:1(d5)/18:0) |
| Cer | Cer(d18:1/22:5) | Cer-d18_1-22_5-264-1 | RP_2 | 4,6 | [M+H]1+ | 612,5 | 264,3 | GlcCer(d18:1(d5)/18:0) |
| HexCer | GlcCer(d18:1/16:0) | HexCer-d18_1-16_0-264-1 | RP_2 | 1,9 | [M+H]1+ | 700,6 | 264,3 | GlcCer(d18:1(d5)/18:0) |
| HexCer | GlcCer(d18:1/16:1) | HexCer-d18_1-16_1-264-1 | RP_2 | 1,8 | [M+H]1+ | 698,6 | 264,3 | GlcCer(d18:1(d5)/18:0) |
| HexCer | GlcCer(d18:1/18:0) | HexCer-d18_1-18_0-264-1 | RP_2 | 2,4 | [M+H]1+ | 728,6 | 264,3 | GlcCer(d18:1(d5)/18:0) |
| HexCer | GlcCer(d18:1/20:0) | HexCer-d18_1-20_0-264-1 | HILIC* | 0,9 | [M+H]1+ | 756,6 | 264,3 | GlcCer(d18:1(d5)/18:0) |
| HexCer | GlcCer(d18:1/22:0) | HexCer-d18_1-22_0-264-1 | RP_2 | 3,8 | [M+H]1+ | 784,7 | 264,3 | GlcCer(d18:1(d5)/18:0) |
| LPC_SIL | PC(18:1(d7)_0:0) | xLPC-18_1-B | HILIC | 2,1 | [M+H]1+ | 529,4 | 184,1 | Stable Isotope Standard |
| LPC | LPC(16:0)_A | LPC-16_0-0-184-1-A | HILIC | 2,1 | [M+H]1+ | 496,3 | 184,1 | PC(18:1(d7)_0:0) |
| LPC | LPC(16:0)_B | LPC-16_0-0-184-1-B | HILIC | 2,2 | [M+H]1+ | 496,3 | 184,1 | PC(18:1(d7)_0:0) |
| LPC | LPC(16:1)_A | LPC-16_1-0-184-1-A | HILIC | 2,1 | [M+H]1+ | 494,3 | 184,1 | PC(18:1(d7)_0:0) |
| LPC | LPC(16:1)_B | LPC-16_1-0-184-1-B | HILIC | 2,2 | [M+H]1+ | 494,3 | 184,1 | PC(18:1(d7)_0:0) |
| LPC | LPC(18:0)_A | LPC-18_0-0-184-1-A | HILIC | 2,0 | [M+H]1+ | 524,4 | 184,1 | PC(18:1(d7)_0:0) |
| LPC | LPC(18:0)_B | LPC-18_0-0-184-1-B | HILIC | 2,1 | [M+H]1+ | 524,4 | 184,1 | PC(18:1(d7)_0:0) |
| LPC | LPC(18:1)_A | LPC-18_1-0-184-1-A | HILIC | 2,0 | [M+H]1+ | 522,4 | 184,1 | PC(18:1(d7)_0:0) |
| LPC | LPC(18:1)_B | LPC-18_1-0-184-1-B | HILIC | 2,1 | [M+H]1+ | 522,4 | 184,1 | PC(18:1(d7)_0:0) |
| LPC | LPC(18:2)_A | LPC-18_2-0-184-1-A | HILIC | 2,1 | [M+H]1+ | 520,3 | 184,1 | PC(18:1(d7)_0:0) |
| LPC | LPC(18:2)_B | LPC-18_2-0-184-1-B | HILIC | 2,1 | [M+H]1+ | 520,3 | 184,1 | PC(18:1(d7)_0:0) |
| PC_SIL | PC(15:0/18:1(d7)) | xPCneg-15_0-18_1-288_1 | RP_1 | 2,2 | [M+CH3COO]1- | 811,6 | 288,3 | Stable Isotope Standard |
| PC | PC(14:0/14:0) | PC-14_0-14_0-227-1 | RP_1 | 1,5 | [M+CH3COO]1- | 736,5 | 227,2 | PC(15:0/18:1(d7)) |
| PC | PC(14:0_16:0) | PC-14_0-16_0-227-1 | RP_1 | 1,9 | [M+CH3COO]1- | 764,5 | 227,2 | PC(15:0/18:1(d7)) |
| PC | PC(14:0_16:1) | PC-14_0-16_1-253-1 | RP_1 | 1,6 | [M+CH3COO]1- | 762,5 | 253,2 | PC(15:0/18:1(d7)) |
| PC | PC(14:0_18:0) | PC-14_0-18_0-227-1 | RP_1 | 2,4 | [M+CH3COO]1- | 792,6 | 227,2 | PC(15:0/18:1(d7)) |
| PC | PC(14:0_18:1) | PC-14_0-18_1-281-1 | RP_1 | 2,0 | [M+CH3COO]1- | 790,6 | 281,2 | PC(15:0/18:1(d7)) |
| PC | PC(14:0_18:2) | PC-14_0-18_2-279-1 | RP_1 | 1,6 | [M+CH3COO]1- | 788,5 | 279,2 | PC(15:0/18:1(d7)) |
| PC | PC(14:1_16:0) | PC-14_1-16_0-225-1 | RP_1 | 1,6 | [M+CH3COO]1- | 762,5 | 225,2 | PC(15:0/18:1(d7)) |
| PC | PC(15:0_16:0) | PC-15_0-16_0-241-1 | RP_1 | 2,1 | [M+CH3COO]1- | 778,6 | 241,2 | PC(15:0/18:1(d7)) |
| PC | PC(15:0_18:1) | PC-15_0-18_1-241-1 | RP_1 | 2,2 | [M+CH3COO]1- | 804,6 | 241,2 | PC(15:0/18:1(d7)) |
| PC | PC(16:0/16:0) | PC-16_0-16_0-255-1 | RP_1 | 2,3 | [M+CH3COO]1- | 792,6 | 255,2 | PC(15:0/18:1(d7)) |
| PC | PC(16:0_16:1) | PC-16_0-16_1-253-1 | RP_1 | 2,0 | [M+CH3COO]1- | 790,6 | 253,2 | PC(15:0/18:1(d7)) |
| PC | PC(16:0_17:0) | PC-16_0-17_0-269-1 | RP_1 | 2,6 | [M+CH3COO]1- | 806,6 | 269,2 | PC(15:0/18:1(d7)) |
| PC | PC(16:0_17:1) | PC-16_0-17_1-267-1-A | RP_1 | 2,2 | [M+CH3COO]1- | 804,6 | 267,2 | PC(15:0/18:1(d7)) |
| PC | PC(16:0_18:0) | PC-16_0-18_0-283-1 | RP_1 | 2,9 | [M+CH3COO]1- | 820,6 | 283,3 | PC(15:0/18:1(d7)) |
| PC | PC(16:0_18:1) | PC-16_0-18_1-281-1 | RP_1 | 2,5 | [M+CH3COO]1- | 818,6 | 281,2 | PC(15:0/18:1(d7)) |
| PC | PC(16:0_18:2) | PC-16_0-18_2-255-1 | RP_1 | 2,1 | [M+CH3COO]1- | 816,6 | 255,2 | PC(15:0/18:1(d7)) |
| PC | PC(16:0_20:4) | PC-16_0-20_4-303-1 | RP_1 | 2,0 | [M+CH3COO]1- | 840,6 | 303,2 | PC(15:0/18:1(d7)) |
| PC | PC(16:0_20:5) | PC-16_0-20_5-301-1 | RP_1 | 1,7 | [M+CH3COO]1- | 838,6 | 301,2 | PC(15:0/18:1(d7)) |
| PC | PC(16:0_22:6) | PC-16_0-22_6-327-1 | RP_1 | 1,9 | [M+CH3COO]1- | 864,6 | 327,2 | PC(15:0/18:1(d7)) |
| PC | PC(16:0_9:0;O) | PC-16_0-9_0al-171-1 | RP_1 | 0,7 | [M+CH3COO]1- | 708,4 | 171,0 | PC(15:0/18:1(d7)) |
| PC | PC(O-16:0/16:0) | PC-16_0e-16_0-255-1 | RP_1 | 2,7 | [M+CH3COO]1- | 778,6 | 255,2 | PC(15:0/18:1(d7)) |
| PC | PC(P-16:0/16:0) | PC-16_0p-16_0-255-1 | RP_1 | 2,6 | [M+CH3COO]1- | 776,6 | 255,2 | PC(15:0/18:1(d7)) |
| PC | PC(16:1_16:1) | PC-16_1-16_1-253-1 | RP_1 | 1,7 | [M+CH3COO]1- | 788,5 | 253,2 | PC(15:0/18:1(d7)) |
| PC | PC(16:1_17:0) | PC-16_1-17_0-253-1 | RP_1 | 2,2 | [M+CH3COO]1- | 804,6 | 253,2 | PC(15:0/18:1(d7)) |
| PC | PC(16:1_18:0) | PC-16_1-18_0-253-1 | RP_1 | 2,5 | [M+CH3COO]1- | 818,6 | 253,2 | PC(15:0/18:1(d7)) |
| PC | PC(16:1_18:1) | PC-16_1-18_1-253-1 | RP_1 | 2,0 | [M+CH3COO]1- | 816,6 | 253,2 | PC(15:0/18:1(d7)) |
| PC | PC(16:1_18:2) | PC-16_1-18_2-279-1 | RP_1 | 1,7 | [M+CH3COO]1- | 814,6 | 279,2 | PC(15:0/18:1(d7)) |
| PC | PC(18:0_18:1) | PC-18_0-18_1-283-1 | RP_1 | 3,1 | [M+CH3COO]1- | 846,6 | 283,3 | PC(15:0/18:1(d7)) |
| PC | PC(18:0_18:2) | PC-18_0-18_2-279-1 | RP_1 | 2,6 | [M+CH3COO]1- | 844,6 | 279,2 | PC(15:0/18:1(d7)) |
| PC | PC(18:1_20:4) | PC-18_0-20_4-303-1 | RP_1 | 2,5 | [M+CH3COO]1- | 868,6 | 303,2 | PC(15:0/18:1(d7)) |
| PC | PC(18:0_9:0;O) | PC-18_0-9_0al-171-1 | RP_1 | 0,8 | [M+CH3COO]1- | 736,4 | 171,0 | PC(15:0/18:1(d7)) |
| PC | PC(18:1/18:1) | PC-18_1-18_1-281-1 | RP_1 | 2,5 | [M+CH3COO]1- | 844,6 | 281,2 | PC(15:0/18:1(d7)) |
| PC | PC(18:1_18:2) | PC-18_1-18_2-279-1 | RP_1 | 2,1 | [M+CH3COO]1- | 842,6 | 279,2 | PC(15:0/18:1(d7)) |
| PC | PC(18:0_20:4) | PC-18_1-20_4-303-1 | RP_1 | 2,1 | [M+CH3COO]1- | 866,6 | 303,2 | PC(15:0/18:1(d7)) |
| PC | PC(18:2/18:2) | PC-18_2-18_2-279-1 | RP_1 | 1,8 | [M+CH3COO]1- | 840,6 | 279,2 | PC(15:0/18:1(d7)) |
| PE_SIL | PE(15:0/18:1(d7) | xPE-15_0-18_1-288_1 | HILIC | 1,6 | [M-H]1- | 709,6 | 288,3 | Stable Isotope Standard |
| PE | PE(14:0_18:1) | PE-14_0-18_1-281-1 | HILIC | 1,6 | [M-H]1- | 688,5 | 281,2 | PE(15:0/18:1(d7) |
| PE | PE(16:0_16:1) | PE-16_0-16_1-253-1 | HILIC | 1,6 | [M-H]1- | 688,5 | 253,2 | PE(15:0/18:1(d7) |
| PE | PE(16:0_18:1) | PE-16_0-18_1-281-1 | HILIC | 1,6 | [M-H]1- | 716,5 | 281,2 | PE(15:0/18:1(d7) |
| PE | PE(16:0_18:2) | PE-16_0-18_2-279-1 | HILIC | 1,6 | [M-H]1- | 714,5 | 279,2 | PE(15:0/18:1(d7) |
| PE | PE(16:0_20:4) | PE-16_0-20_4-303-1 | HILIC | 1,6 | [M-H]1- | 738,5 | 303,2 | PE(15:0/18:1(d7) |
| PE | PE(16:1_18:0) | PE-16_1-18_0-283-1 | HILIC | 1,6 | [M-H]1- | 716,5 | 283,3 | PE(15:0/18:1(d7) |
| PE | PE(16:1_18:1) | PE-16_1-18_1-253-1 | HILIC | 1,6 | [M-H]1- | 714,5 | 253,2 | PE(15:0/18:1(d7) |
| PE | PE(18:0_18:1) | PE-18_0-18_1-281-1 | HILIC | 1,6 | [M-H]1- | 744,6 | 281,2 | PE(15:0/18:1(d7) |
| PE | PE(18:0_18:2) | PE-18_0-18_2-279-1 | HILIC | 1,6 | [M-H]1- | 742,5 | 279,2 | PE(15:0/18:1(d7) |
| PE | PE(18:0_20:4) | PE-18_0-20_4-303-1 | HILIC | 1,6 | [M-H]1- | 766,5 | 303,2 | PE(15:0/18:1(d7) |
| PE | PE(18:1/18:1) | PE-18_1-18_1-281-1 | HILIC | 1,6 | [M-H]1- | 742,5 | 281,2 | PE(15:0/18:1(d7) |
| PE | PE(18:1_18:2) | PE-18_1-18_2-279-1 | HILIC | 1,6 | [M-H]1- | 740,5 | 279,2 | PE(15:0/18:1(d7) |
| PE | PE(18:1_20:4) | PE-18_1-20_4-303-1 | HILIC | 1,6 | [M-H]1- | 764,5 | 303,2 | PE(15:0/18:1(d7) |
| LPE_SIL | PE(18:1(d7)_0:0) | xLPE-18_1-A | HILIC | 2,2 | [M-H]1- | 487,3 | 346,3 | Stable Isotope Standard |
| PEO | PE (O-2:0_18:1) | PEO-2_0-18_1-367-1 | HILIC | 1,9 | [M-H]1- | 508,3 | 367,3 | PE(18:1(d7)_0:0) |
| PEO | PE (O-2:0_18:2) | PEO-2_0-18_2-365-1 | HILIC | 1,9 | [M-H]1- | 506,3 | 365,3 | PE(18:1(d7)_0:0) |
| PEO | PE (O-2:0_20:1) | PEO-2_0-20_1-395-1 | HILIC | 1,9 | [M-H]1- | 536,4 | 395,4 | PE(18:1(d7)_0:0) |
| PEO | PE (O-2:0_20:4) | PEO-2_0-20_4-389-1 | HILIC | 1,9 | [M-H]1- | 530,3 | 389,3 | PE(18:1(d7)_0:0) |
| PG_SIL | PG(15:0/18:1(d7)) | xPG-15_0-18_1-288_1 | HILIC | 1,2 | [M-H]1- | 740,6 | 288,3 | Stable Isotope Standard |
| PG | PG(14:0_16:0) | PG-14_0-16_0-227-1 | HILIC | 1,3 | [M-H]1- | 693,5 | 227,2 | PG(15:0/18:1(d7)) |
| PG | PG(16:0/16:0) | PG-16_0-16_0-255-1 | HILIC | 1,2 | [M-H]1- | 721,5 | 255,2 | PG(15:0/18:1(d7)) |
| PG | PG(16:0_16:1) | PG-16_0-16_1-253-1 | HILIC | 1,2 | [M-H]1- | 719,5 | 253,2 | PG(15:0/18:1(d7)) |
| PG | PG(16:0_18:0) | PG-16_0-18_0-255-1 | HILIC | 1,2 | [M-H]1- | 749,5 | 255,2 | PG(15:0/18:1(d7)) |
| PG | PG(16:0_18:1) | PG-16_0-18_1-281-1 | HILIC | 1,2 | [M-H]1- | 747,5 | 281,2 | PG(15:0/18:1(d7)) |
| PG | PG(16:0_18:2) | PG-16_0-18_2-279-1 | HILIC | 1,2 | [M-H]1- | 745,5 | 279,2 | PG(15:0/18:1(d7)) |
| PG | PG(16:1_18:0) | PG-16_1-18_0-253-1 | HILIC | 1,2 | [M-H]1- | 747,5 | 253,2 | PG(15:0/18:1(d7)) |
| PG | PG(16:1_18_1) | PG-16_1-18_1-253-1 | HILIC | 1,2 | [M-H]1- | 745,5 | 253,2 | PG(15:0/18:1(d7)) |
| PG | PG(18:0_18:1) | PG-18_0-18_1-281-1 | HILIC | 1,2 | [M-H]1- | 775,6 | 281,2 | PG(15:0/18:1(d7)) |
| PG | PG(18:0_18_2) | PG-18_0-18_2-279-1 | HILIC | 1,2 | [M-H]1- | 773,5 | 279,2 | PG(15:0/18:1(d7)) |
| PG | PG(18:0_20:4) | PG-18_0-20_4-303-1 | HILIC | 1,2 | [M-H]1- | 797,5 | 303,2 | PG(15:0/18:1(d7)) |
| PG | PG(18:1_18:1) | PG-18_1-18_1-281-1 | HILIC | 1,2 | [M-H]1- | 773,5 | 281,2 | PG(15:0/18:1(d7)) |
| PG | PG(18:1_18:2) | PG-18_1-18_2-279-1 | HILIC | 1,2 | [M-H]1- | 771,5 | 279,2 | PG(15:0/18:1(d7)) |
| PI | PI(15:0/18:1(d7)) | xPI-15_0-18_1-288_1 | | 2,4 | | 828,6 | 288,3 | Stable Isotope Standard |
| PI | PI(16:0_18:1) | PI-16_0-18_1-255-1 | HILIC | 2,4 | [M-H]1- | 835,5 | 255,2 | xPI-15_0-18_1-288_1 |
| PI | PI(18:0_18:1) | PI-18_0-18_1-283-1 | HILIC | 2,4 | [M-H]1- | 863,6 | 283,3 | xPI-15_0-18_1-288_2 |
| PI | PI(18:0_20:4) | PI-18_0-20_4-283-1 | HILIC | 2,3 | [M-H]1- | 885,6 | 283,3 | xPI-15_0-18_1-288_3 |
| PI | PI(18:1/18:1) | PI-18_1-18_1-281-1 | HILIC | 2,4 | [M-H]1- | 861,6 | 281,2 | xPI-15_0-18_1-288_4 |
| SM d18_1 | SM(d18:1_18:1(d9)) | xSM-d18_1-18_1_1 | HILIC | 1,9 | [M+H]1+ | 738,7 | 184,1 | Stable Isotope Standard |
| SM d18_1 | SM(30:1;O2) | SM d18_1-12_0 | HILIC | 2,0 | [M+H]1+ | 647,5 | 184,1 | xSM-d18_1-18_1_1 |
| SM d18_1 | SM(32:1;O2) | SM d18_1-14_0 | HILIC | 2,0 | [M+H]1+ | 675,5 | 184,1 | xSM-d18_1-18_1_2 |
| SM d18_1 | SM(32:2;O2) | SM d18_1-14_1 | HILIC | 2,0 | [M+H]1+ | 673,5 | 184,1 | xSM-d18_1-18_1_3 |
| SM d18_1 | SM(34:1;O2) | SM d18_1-16_0 | HILIC | 1,9 | [M+H]1+ | 703,6 | 184,1 | xSM-d18_1-18_1_4 |
| SM d18_1 | SM(34:2;O2) | SM d18_1-16_1 | HILIC | 1,9 | [M+H]1+ | 701,6 | 184,1 | xSM-d18_1-18_1_5 |
| SM d18_1 | SM(35:1;O2) | SM d18_1-17_0 | HILIC | 1,9 | [M+H]1+ | 717,6 | 184,1 | xSM-d18_1-18_1_6 |
| SM d18_1 | SM(36:1;O2) | SM d18_1-18_0 | HILIC | 1,9 | [M+H]1+ | 731,6 | 184,1 | xSM-d18_1-18_1_7 |
| SM d18_1 | SM(36:2;O2) | SM d18_1-18_1 | HILIC | 1,9 | [M+H]1+ | 729,6 | 184,1 | xSM-d18_1-18_1_8 |
| SM d18_1 | SM (40:1;O2) | SM d18_1-22_0 | HILIC | 1,9 | [M+H]1+ | 787,7 | 184,1 | xSM-d18_1-18_1_9 |
| SM d18_1 | SM (42:1;O2) | SM d18_1-24_0 | HILIC | 1,8 | [M+H]1+ | 815,7 | 184,1 | xSM-d18_1-18_1_10 |
| SM d18_1 | SM (42:2;O2) | SM d18_1-24_1 | HILIC | 1,8 | [M+H]1+ | 813,7 | 184,1 | xSM-d18_1-18_1_11 |

Lipid structures and masses can be found on lipid maps webpage: https://lipidmaps.org

*GlcCer were run on both HILIC and RP for method comparision. For GlcCer(d18:1/20:0) only data from HILIC method was aquired by mistake and this result was used for quantification.For the other GlcCer the RP results was used. Comparision between HILIC and RP showed good agreement for GlcCer analysed on both methods.

| Mobile Phase | A1 | B1 | Strong Needle Wash | Weak Needle Wash |
|---|---|---|---|---|
| Solvent | ACN:H2O [95:5] | ACN:H2O [50:50] | IPA [100%] | ACN:H2O [95:5] |
| Buffer AmAC | 10mM | 10mM | - | - |

| Time (min) | %A | B% | Flow (ml/min) | Curve |
|---|---|---|---|---|
| 0 | 99,9 | 0,1 | 0,6 | 6 |
| 2 | 80 | 20 | 0,6 | 6 |
| 5 | 20 | 80 | 0,6 | 6 |
| 5,5 | 20 | 80 | 0,6 | 6 |
| 6 | 99,9 | 0,1 | 0,6 | 6 |
| 10 | 99,9 | 0,1 | 0,6 | 6 |

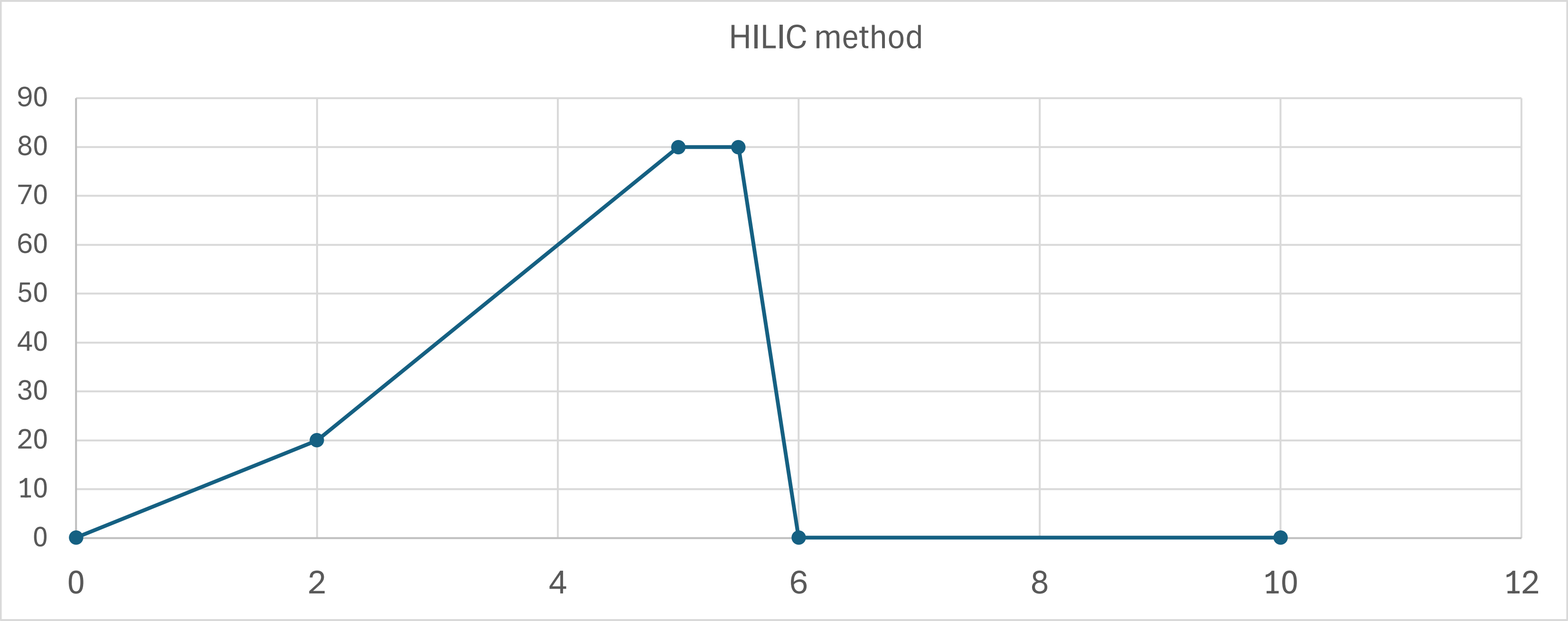

| Mobile Phase | A1 | B1 | Strong Needle Wash | Weak Needle Wash |
|---|---|---|---|---|
| Solvent | H2O:ACN [90:10] | ACN:IPA:H2O [70:20:10] | ACN:IPA:H2O [70:20:10] | H20:ACN:IPA:MeOH [1:1:1:1] |
| Buffer AmAc | 10mM | 10mM | - | - |

**RP_1**

| Row | Time (min) | %A | B% | Flow (ml/min) | Curve |
|---|---|---|---|---|---|
| 1 | 0 | 9 | 91 | 0,6 | initial |
| 2 | 4 | 2,2 | 97,8 | 0,6 | 6 |
| 3 | 4,01 | 0 | 100 | 0,6 | 6 |
| 4 | 5,5 | 0 | 100 | 0,6 | 6 |
| 5 | 5,51 | 9 | 91 | 0,6 | 6 |
| 6 | 7 | 9 | 91 | 0,6 | 6 |

**RP_2**

| Row | Time (min) | %A | B% | Flow (ml/min) | Curve |
|---|---|---|---|---|---|
| 1 | 0 | 9,0 | 91,0 | 0,6 | initial |
| 2 | 4 | 2,2 | 97,8 | 0,6 | 6 |
| 3 | 4,01 | 0,1 | 99,9 | 0,6 | 6 |
| 4 | 7,90 | 0,1 | 99,9 | 0,6 | 6 |
| 5 | 8 | 9,0 | 91,0 | 0,6 | 6 |
| 6 | 11 | 9,0 | 91,0 | 0,6 | 6 |

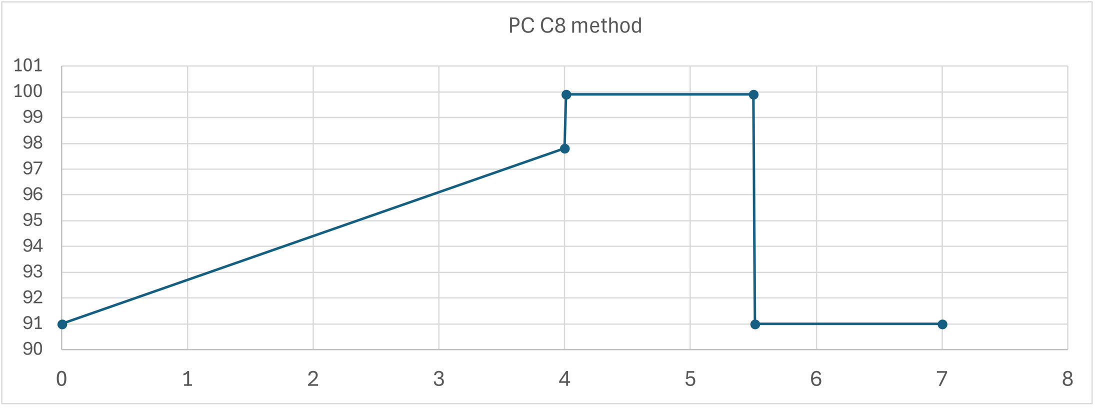


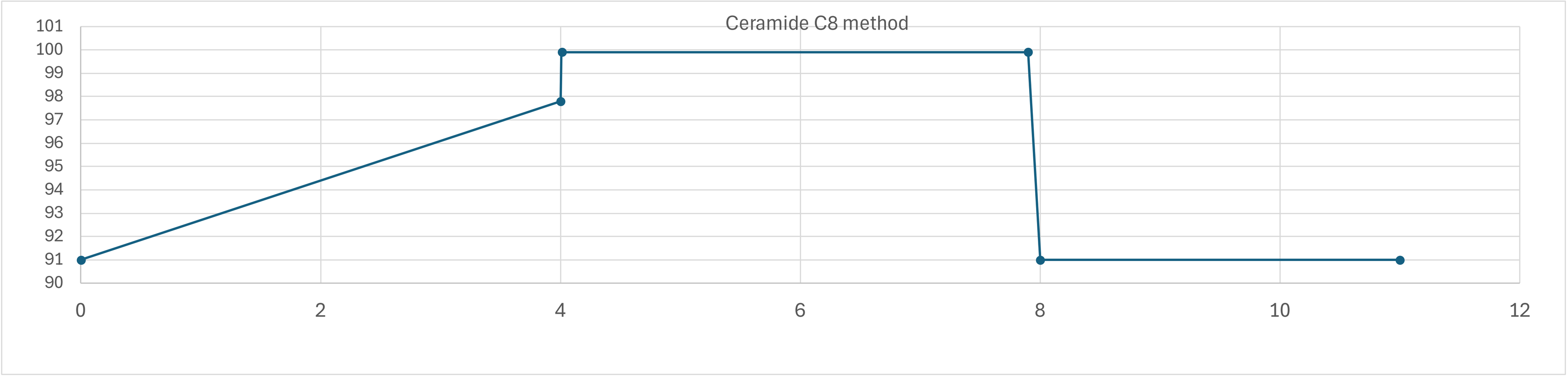

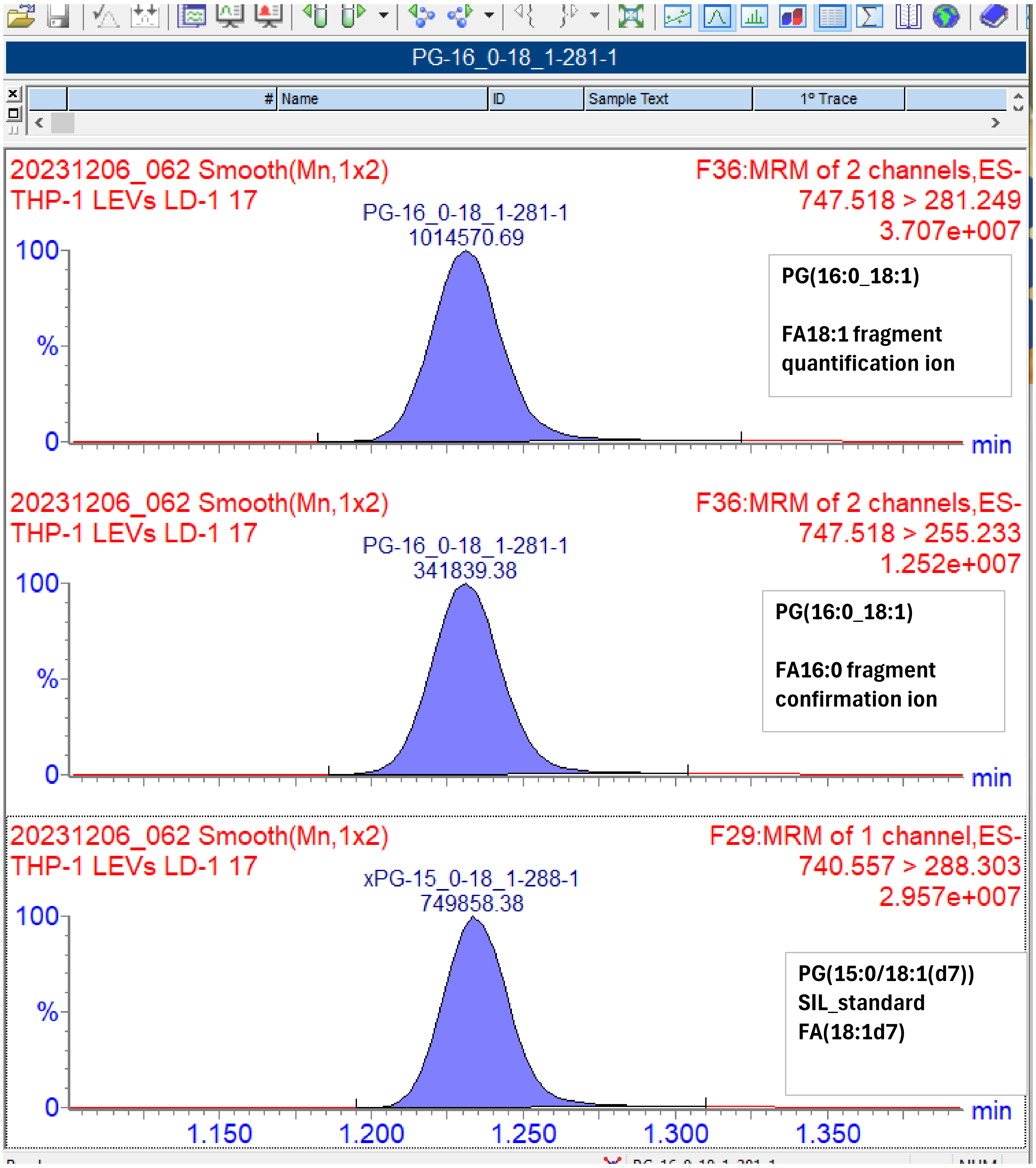

PG-16_0-18_1-281-1
### Name
ID
Sample Text
1° Trace
20231206_062 Smooth(Mn,1x2)
THP-1 LEVs LD-1 17
F36:MRM of 2 channels,ES-
747.518 > 281.249
3.707e+007
PG-16_0-18_1-281-1
1014570.69
PG(16:0_18:1)
FA18:1 fragment quantification ion
20231206_062 Smooth(Mn,1x2)
THP-1 LEVs LD-1 17
F36:MRM of 2 channels,ES-
747.518 > 255.233
1.252e+007
PG-16_0-18_1-281-1
341839.38
PG(16:0_18:1)
FA16:0 fragment confirmation ion
20231206_062 Smooth(Mn,1x2)
THP-1 LEVs LD-1 17
F29:MRM of 1 channel,ES-
740.557 > 288.303
2.957e+007
xPG-15_0-18_1-288-1
749858.38
PG(15:0/18:1(d7))
SIL_standard
FA(18:1d7)
100
%
0
min
1.150
1.200
1.250
1.300
1.350
